# A global map of human pressures on tropical coral reefs

**DOI:** 10.1101/2021.04.03.438313

**Authors:** Marco Andrello, Emily Darling, Amelia Wenger, Andrés F. Suárez-Castro, Sharla Gelfand, Gabby N. Ahmadia

## Abstract

As human activities on the world’s oceans intensify, mapping human pressures is essential to develop appropriate conservation strategies and prioritize investments with limited resources. Here, we map non-climatic pressures on coral reefs using the latest quantitative data layers on fishing, nitrogen and sediment pollution, coastal and industrial development, and tourism. Across 54,596 coral reef pixels worldwide, we identify the top-ranked local pressure and estimate a cumulative pressure index mapped at 0.05-degree (∼5 km) resolution. Fishing was the most common top-ranked pressure followed by water pollution (nutrients and sediments), although there is substantial variation by regions. We also find that local pressures are similar inside and outside a proposed global portfolio of coral reef climate refugia. We provide the best available information to inform critical conservation strategies and ensure local pressures are effectively managed to increase the likelihood of the persistence of coral reefs to climate change.

## Introduction

Marine biodiversity globally is threatened by the expanding and intensifying impacts of human activities and climate change (O’Hara, Frazier, & Halpern 2021). Implementing solutions to mitigate pressures on ocean ecosystems is a key strategy to slow and reverse biodiversity decline and maintain ecosystem functioning and integrity. Being more intentional about how we allocate critical but limited conservation resources to manage pressures will be critical to achieve the Convention on Biological Diversity’s Post-2020 Global Biodiversity Framework. Mapping pressures on ecosystems can predict ecological conditions to prioritize climate adaptation or mitigation responses (Grantham et al., 2020) and identify the most threatening activities to ecological resilience and integrity (Tulloch et al., 2015). Failing to address the most impactful pressures can undermine the effectiveness of conservation interventions, so tailored approaches are needed to guide appropriate conservation strategies at local scales and avoid further ecosystem degradation (Allan et al., 2019; Tulloch et al., 2020).

Coral reefs are among the most diverse marine ecosystems on the planet, a critical source of livelihoods, cultural and food security for millions of people and are declining at alarming rates from global and local pressures (Eddy, Cheung, & Bruno, 2018). As impacts of thermal stress, coral bleaching, ocean acidification, and sea level rise intensify (Hoegh-Guldberg et al., 2018), decarbonization is a defining challenge for the survival of functioning coral reefs (Darling et al., 2019; Morrison et al., 2019). Complementary to decarbonization, addressing non-climatic pressures remains crucial for maintaining coral reef function, and is a foundation of coral reef conservation initiatives worldwide. Fishing, pollution, and development can affect coral reefs worldwide (with the exception of some remote ‘wilderness’ areas) and can have damaging population and ecosystem-level effects (Burke, Reytar, Spalding, & Perry, 2011). Identifying and addressing non-climatic pressures can enhance recovery from climate shocks (Claar et al., 2020), reduce coral disease (Lamb et al., 2016), and is particularly important within potential climate refugia that are hoped to maintain functioning coral reefs over the coming decades (Côté, Darling, & Brown, 2016; Beyer et al. 2018; Darling et al., 2019).

One of the first global assessments of human pressures on coral reefs was The Reefs at Risk project that mapped the impacts of overfishing and destructive fishing, sediment and nitrogen pollution, marine-based pollution and damage, coastal development, thermal stress and ocean acidification on coral reefs (Burke, Bryant, McManus, & Spalding, 1998; Burke et al., 2011). Reefs at Risk has been a valuable tool guiding conservationists and researchers for the past twenty years. However, recent advances in satellite imagery and analytical approaches have improved our ability to map human pressures on coral reefs. These advances include a global ‘gravity’ index of fishing pressure based on travel time to markets (Cinner et al., 2016, 2018), more realistic data layers to model soil erosion and sediment runoff (Borrelli et al., 2017), and incorporating more recent and highly resolved estimates of humans populations (Center for International Earth Science Information Network - CIESIN -Columbia University, 2018).

Here, we provide an updated global mapping of local pressures on the world’s coral reefs, and provide quantitative data layers of human pressures to coral reefs. This advances the ‘low’, ‘medium’, ‘high’ pressure categories of the Reefs and Risk approach, and we make the underlying datasets and a series of report cards freely available (see Acknowledgements and Data). Our analyses identify reef areas with the highest impact for each pressure independently and cumulative pressures, and determine the top-ranked pressure. We also evaluated local pressures across a global portfolio of proposed climate refugia for coral reefs (Beyer et al., 2018), which can inform the conservation efforts of several multi-million dollar global conservation initiatives, including Bloomberg Philathropies Vibrant Oceans Initiative, the Coral Reef Rescue Initiative, and the United Nations’ Global Coral Reef Fund. Intended to be combined with local knowledge and expertise, our analyses will support decision makers to identify and mitigate the top pressures on coral reefs to ensure the persistence of coral reefs and their associated wealth of services to humanity.

## Methods

### Data layers

Tropical coral reef locations were used from Beyer et al. (2018) based on the Global Distribution of Coral Reefs dataset (UNEP-WCMC, WorldFish Centre, WRI, & TNC, 2010) and updates from the National Oceanic and Atmospheric Administration (NOAA). We mapped reef locations onto a 0.05-degree resolution (∼5 km) raster dataset, resulting in 54,596 reef-containing raster grid cells worldwide.

We integrated spatial data layers on six local pressures on coral reefs: fishing, coastal development, industrial development, tourism, sediment pollution and nitrogen pollution (**Table 1**). We define a ‘pressure’ as a contextual variable that can, under certain conditions or above a specific magnitude, exert unsustainable pressure that degrades the ecological function, productivity, or resilience of coral reef ecosystems. While pressures can result in negative consequences for reefs, they are also crucial activities for livelihoods and wellbeing (e.g., small-scale fisheries and tourism); the goal of our analysis is thus to inform priorities for sustainable management that can ensure the long term persistence of coral reef ecosystems and coastal communities who depend on reef-related ecosystem services. More details about the definition and calculation of each pressure can be found in the **Supplementary Methods**.

**Table 1.** Data layers used to quantify six local pressures on coral reef biodiversity, functioning and persistence.

| Data layer | Source | Description | Original resolution | Perceived impact score from Wear (2016)<sup>1</sup> |
| --- | --- | --- | --- | --- |
| Fishing (market gravity) | Cinner et al. (2016, 2018) | From economic geography, ‘gravity’ evaluates the potential interactions between human populations and coral reefs accounting for both population size and accessibility (travel time) to coral reefs. Market gravity characterizes the market-driven influences from major population centers on coral reef fish biomass and fisheries. High market gravity, measured as (number of people) / (hours of travel)<sup>2</sup> has been shown to reduce fish biomass and the occurrence of top predators on coral reefs. | 10 km grid cells | Overfishing (4.3) |
| Coastal development | Gridded Population of the World, v4 (Center for International Earth Science Information Network - CIESIN - Columbia University, 2018) | Coastal development can affect nearshore ecosystems through wastewater discharge (Wear & Thurber, 2015), construction (Wenger et al., 2017), and land-based agriculture and industry (Wenger et al., 2020). As an indicator of coastal development, we used the number of people living within a 5km buffer of each coral reef cell using a global data layer of 2020 human populations. | 2.5 degree minutes (~5 km) | Coastal development (4.3) |
| Industrial development | World Ports, point locations of existing ports, freely available from Google data (https://goo.gl/Yu8xxt). | We used port locations as a proxy for potential pressures from industrial development, including maritime activities such as dredging (Yap & Lam, 2013). The dataset identifies 2,646 coastal ports, and we counted all ports within 5 km<sup>2</sup> as this is the maximum likely distance of dredging impacts (Wenger et al., 2018). | Point locations | Coastal development (4.3) |
| Tourism | Spalding et al. (2017) | An estimate of the annual economic value of coral reefs to the tourism sector, including recreational diving and snorkelling, the provision of calm waters, coral sand beaches, views and seafood. Economic value (in 2013 U.S. dollars) is estimated for coral reefs globally with tourism activity based on data from ~2005-2012. | 0.5 km grid cells | Coastal development (4.3) |
| Water pollution (sediments) | This paper | We combined two approaches for estimating sediment delivery to coral reefs (see Supplementary Methods). Sediment exposure incorporates land cover and management, rainfall-runoff erosivity, slope, steepness, and soil erodibility. Sediment exposure to adjacent coral reefs was then modelled using a sediment plume model, and estimated as tons of sediment / km<sup>2</sup>. | 1 km raster grid | Watershed pollution (3.8) |
| Water pollution (nitrogen) | This paper | Nitrogen delivery to coral reefs is based on catchment-level crop cover and national nitrogen fertilizer use reported by the Food and Agriculture Organization (FAO) and nitrogen plume models, as detailed in the Supplementary Methods. | 1 km raster grid | Watershed pollution (3.8) |

<sup>1</sup>Weights are estimated on a scale from 0 (low) to 6 (high).

To compare local pressures inside and outside potential climate refugia, we used the Beyer et al. (2018) ‘50 Reefs’ analysis which identified 19,161 coral reef cells (out of 54,596, or 35.1%) as potential climate refugia based on an analysis of past, present, and future climate, cyclones and larval connectivity. These cells are grouped into 83 bioclimatic units (BCUs), each containing approximately 500 km^2^ of coral reef habitat located over a spatially contiguous area.

### Analysis

We first converted a raster dataset of coral reef locations into a vector layer of 54,596 square reef polygons (0.05° x 0.05°; approximately 5 km x 5 km) and projected each pressure data layer (**Table 1**) onto this dataset using a WGS84 coordinate reference system. To rank pressures within each cell and minimize the impact of extreme values, we calculated the percentile of each pressure within a cell from the global distribution of the pressure (see **Supplementary Methods)**. The top pressure within each cell was identified as the pressure with the highest percentile compared to other pressures.

A cumulative pressure impact score for each reef pixel was estimated from the weighted average of the percentiles of the six pressure layers. Weights were determined from Wear (2016) from a survey of 170 managers in 50 coral reef countries and territories to calculate the perceived severity of different pressures on coral reefs (**Table 1**).

To evaluate pressures within potential climate refugia (BCUs), we extracted the median percentile of reef cells within a BCU for each pressure and compared the median percentiles across pressure data layers; the top-ranked pressure within a BCU was the pressure with the highest median percentile. However, the spatial variability of pressures within each BCU is also important for conservation and management planning (see **Box 1**).

## Results

### Ranking local pressures

Fishing (market gravity) was the most frequently top-ranked local pressure in 16,543 out of 54,596 reef cells, or 30.3% of coral reefs worldwide (**Figure 1**). Coastal development was the second most common top-ranked pressure (10,630 reef cells, or 19.5% of all cells), followed by industrial development (1,425, or 2.6%), tourism (8,386, or 15.4%), sediment pollution (9,046, or 16.6%) and nitrogen pollution (8,566, or 15.7%). Combining the two water pollution pressures (nitrogen and sediment) raises water pollution to the most common global pressure in 17,612 reef cells (or 32.2%). In total, fishing and water pollution are the top-ranked pressures in 34,155 reef cells, or 62.6% of the world’s coral reefs.

**Figure 1.**
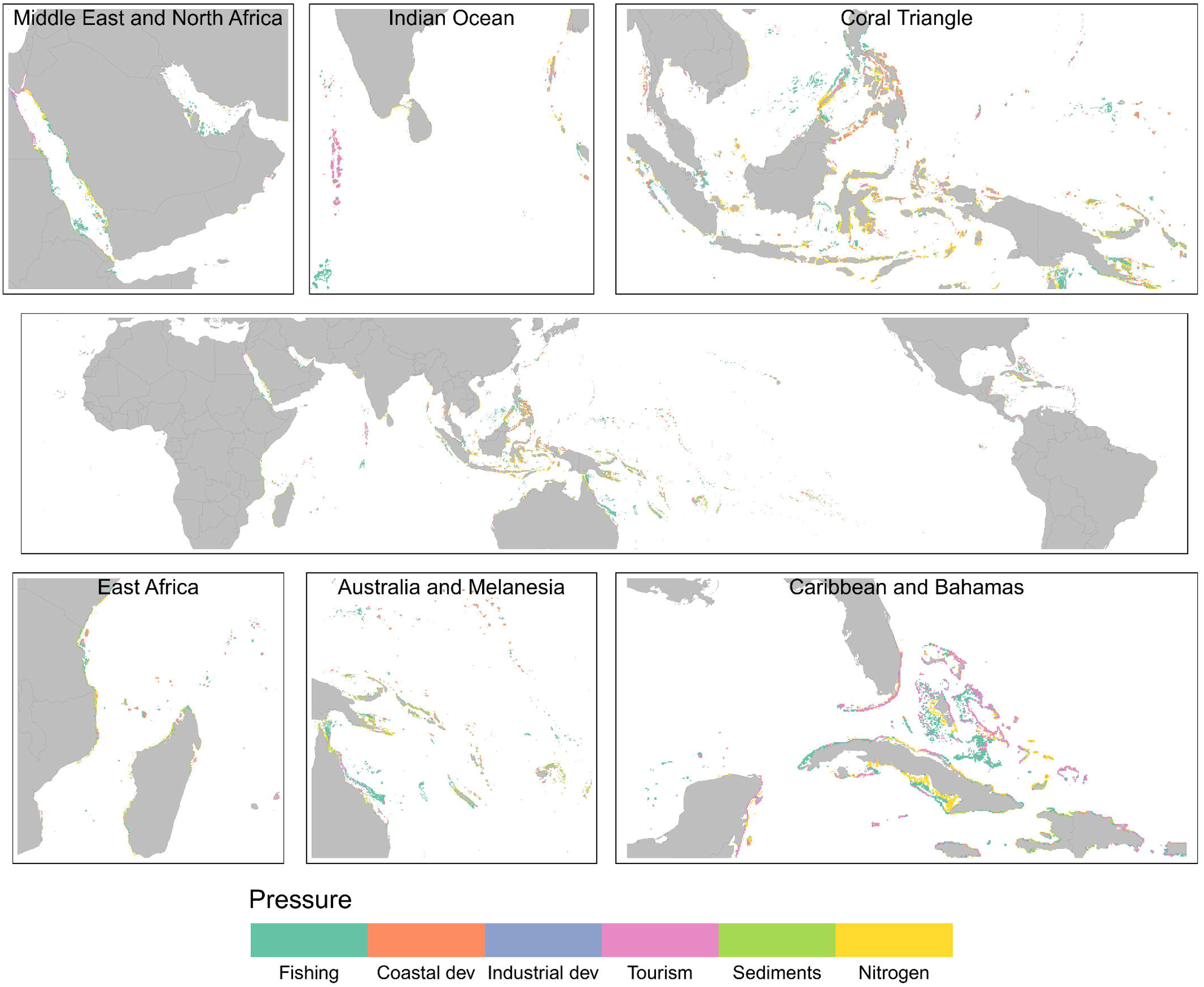
Top-ranked local pressures for coral reefs. The top-ranked pressure in each coral reef cell was identified by comparing each cell’s percentiles across different pressure layers. Colours indicate the top ranked pressure. Panels show global (middle) and insets of key coral reef geographies (top and bottom panels).

Comparing the intensity and spatial distribution of individual local pressures shows that fishing and coastal development are diffuse pressures affecting reefs at large spatial scales while the influences of tourism, sediment and nitrogen pollution are more spatially clustered, and the effects of industrial development are visible only at very local scales (**Figures S1-S6)**.

All coral reef regions have reefs with high cumulative impact scores, which are often clustered into contiguous areas distributed over the entire geographical range (**Figure 2**). Areas with higher cumulative impact scores are typically located near the coasts of continents and islands, while remote reefs, not surprisingly, have lower cumulative impact scores.

**Figure 2.**
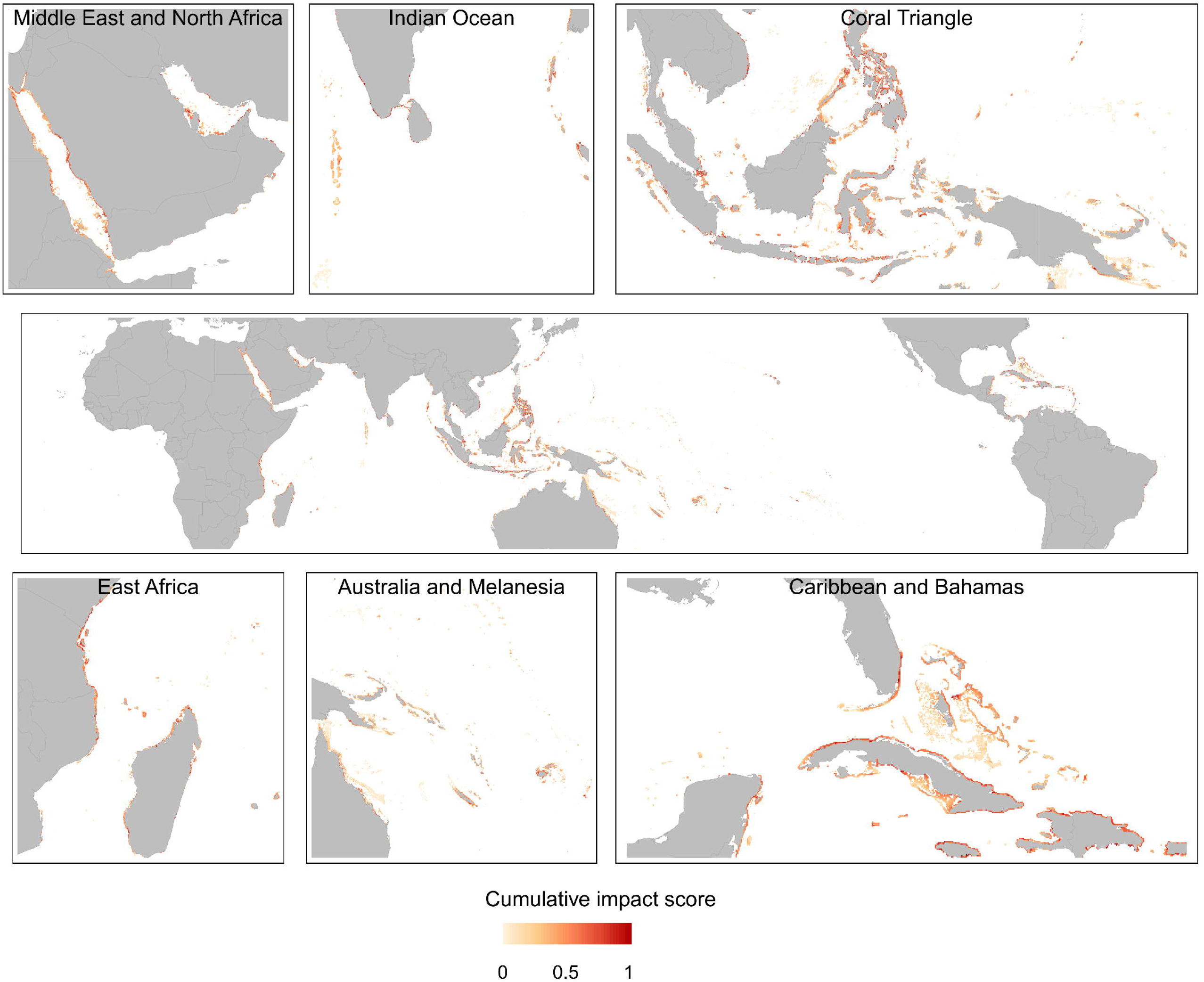
Cumulative impact of local pressures on coral reefs. Cumulative impact scores for each reef cell, estimated by combining pressure scores of each of the different data layers in Figure 1. Panels show global (middle) and insets of key coral reef geographies (top and bottom panels); darker colours indicate higher cumulative pressure scores than lighter colours (lower cumulative scores).

### Regional variation

There was substantial variation within- and between-regions in individual and cumulative pressures (**Figure 3;** see **Figure S11** for region classifications). The Western Indian Ocean, Southeast Asia and North Pacific Ocean regions had high, on average, pressures compared to other regions. Within-region variation was also notable. For example, while the Central Indian Ocean region has low regional averages for sediment and nitrogen pollution pressures, some reefs in this region have the highest values (percentiles) of these pressures globally (e.g., high sedimentation from upstream logging activities in Madagascar; Maina et al. 2013) (see outliers in **Figure 3**). Similarly, while Pacific regions (Australia, Micronesia, Polynesia and Melanesia) have the lowest median fishing pressures by region, individual reef cells within these regions have some of highest fishing pressure percentiles globally. Regional variability further highlights the need to match conservation and management interventions to the appropriate scale and location of local pressures.

**Figure 3.**
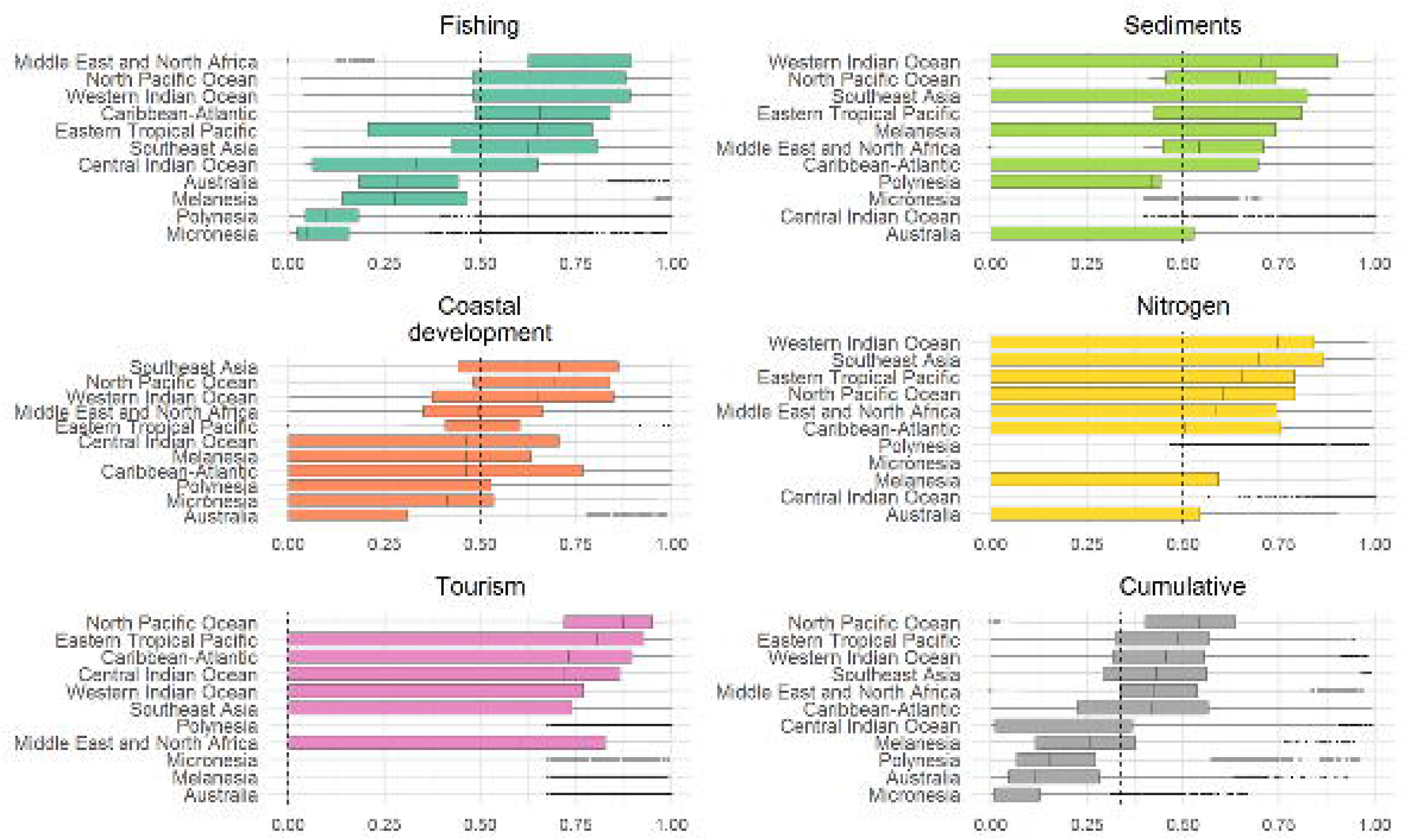
Regional comparisons of individual and cumulative local pressures on coral reefs. Dashed line shows global median, and boxplots show the 25th, 50th (median) and 75th quantile; outliers are points beyond the whiskers (1.5 * interquartile range, or the distance between the first and third quartiles). pressure values (x-axis) are distributed from 0 (lowest value in data layer) to 1 (highest value in data layer).

We also observed variation in the relative distribution of top-ranked pressures by region (**Figure S7**). Fishing was a top-ranked pressure in all regions, but in different proportions relative to other top-ranked pressures, ranging from 12.3% of reef cells in the Eastern Tropical Pacific to 49.0% in Australia. Water pollution (sediments and nitrogen) exhibited even larger regional variation, from 1.4% of reef cells in Micronesia to 46.8% of reef cells in Melanesia. Interestingly, every region had one of the six pressures ranked as a top pressure in some reef cells (**Figure S7**).

Top-ranked pressures were typically associated with high percentiles of the pressure, but there were also cells where top-ranked threats comprised low percentiles, especially for fishing (**Figure S8**). These cells were typically located in remote areas where fishing and other pressures are likely limited, for example the uninhabited islands and reefs of the Chagos Archipelago, the Federated States of Micronesia, New Caledonia and Tuvalu, and more remote islands of Australia’s Great Barrier Reef (**Figure S1**). In contrast, industrial development was ranked as a top-pressure only at very high levels of impact (**Figure S3**).

### Climate refugia

The identity and magnitude of local pressures were fairly consistent inside and outside a proposed global portfolio of climate refugia (Beyer et al. 2018) (**Figure 4**). Reef cells inside and outside bioclimatic units (BCUs, or potential climate refugia) were exposed to similar levels of non-climate pressures, although reef cells inside BCUs consistently had more cells with higher impacts of local pressures than non-BCU cells (**Figure 4)**.

**Figure 4.**
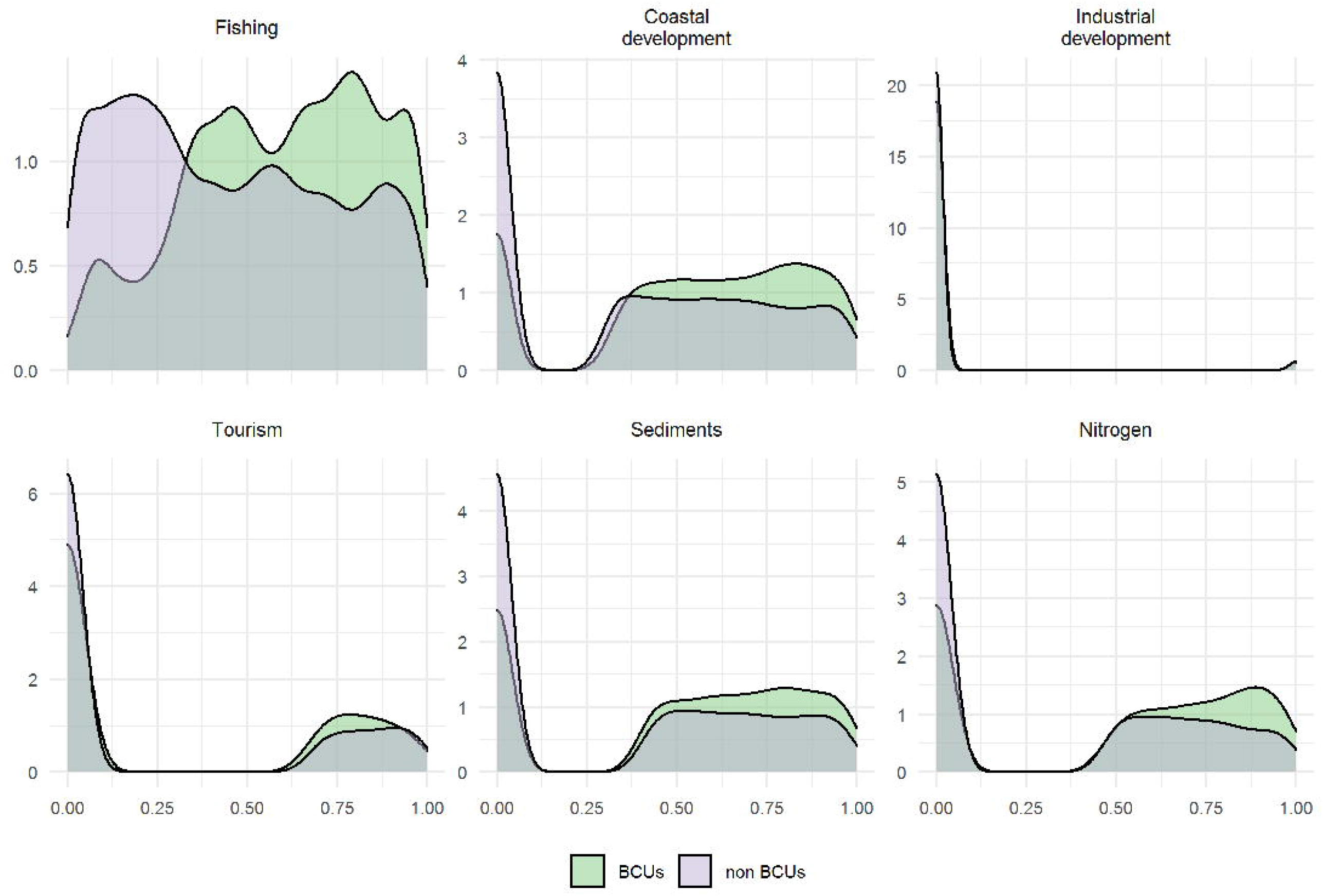
Local pressures across climate refugia. Density distribution of local pressure values inside a global portfolio of potential climate refugia (BCU reef cells, n = 19,161) compared to sites expected to have higher climate impacts (non-BCU reef cells, *n* = 35,435). pressure values range from 0 (lowest value in global data layer) to 1 (highest value) and the y-axis shows the frequency of values.

Fishing was identified as the top pressure in 27 of the 83 BCUs (33%), coastal development in 15 BCUs (18%), tourism in 10 BCUs (12%), sediment pollution in 12 BCUs (14%) and nitrogen pollution in 19 BCUs (23%); industrial development is not identified a top pressure for any BCU (**Figure 5**). BCUs in Southeast Asia, Middle East and North Africa, East Africa, and the Caribbean-Atlantic have high cumulative pressure scores, suggesting these potential climate refugia face substantial human pressures that require appropriate and effective conservation and management interventions. Conversely, BCUs in Australia, Micronesia and Polynesia often have median pressure percentiles and median cumulative impact scores below the global median (**Figure 5**). The top-ranked pressures were similar between BCU and non-BCU reef cells (**Figure S10**).

**Figure 5.**
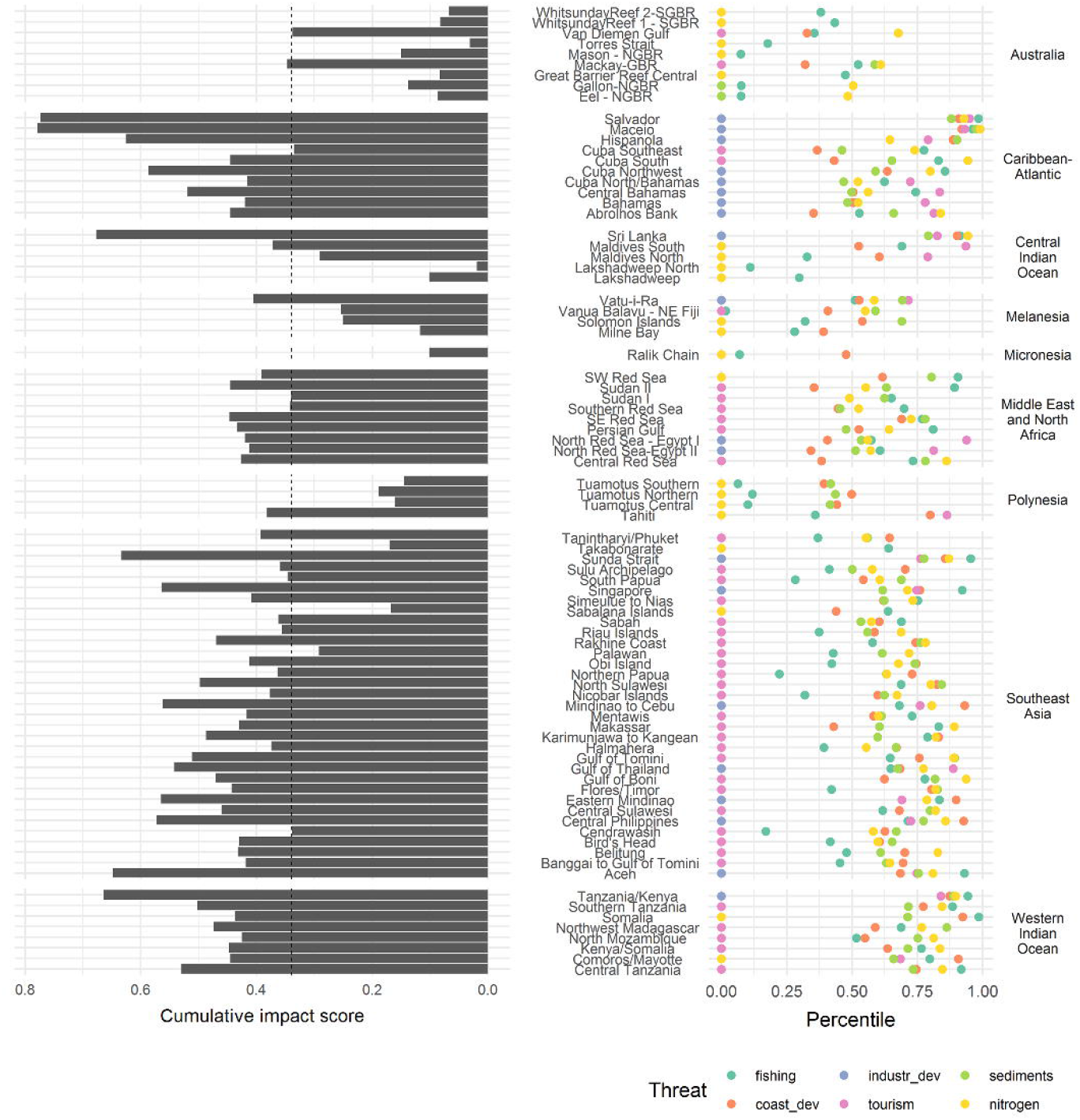
Top-ranked and cumulative non-climate pressures on a global portfolio of potential climate refugia. Across nine regions, BCUs occur across a gradient of cumulative impact (left panel; vertical dashed line is the median cumulative impact score of the 54,595 reef cells worldwide). The right panel shows the distribution of median pressure percentiles for the reef cells within each bioclimatic unit (BCU; Beyer et al. 2018). pressure percentiles (x-axis) range from 0 (lowest value in pressure data layer) to 1 (highest value). The top-ranked pressure is identified as the pressure with the highest percentile (median of all reef cells within a BCU).

## Discussion

Achieving conservation outcomes requires identifying top pressures and applying appropriate management interventions to reduce the impact of unsustainable pressures (Tulloch et al. 2015). Here, we map local pressures at a high-resolution across 54,596 coral reef pixels globally. Our findings improve past global assessments to map pressures on coral reefs. In the Reefs at Risk project (Burke et al., 2011), overfishing and coastal development were identified as the two top pressures affecting about half of coral reefs globally. Our findings suggest that sediment and nutrient pollution are the main impacts of coastal development affecting coral reefs, suggesting integrated watershed management to improve water quality is a key component for sustainable coastal development. A subsequent survey of managers (Wear 2016) confirmed the results of the Reefs at Risk analysis but added watershed-based pollution as a top pressure. We find similar results to both studies, with fishing and water pollution (sediment and nitrogen delivery) identified as the top two pressures on coral reefs affecting 34,155 out of 54,596 reef cells (62.6% of the world’s coral reefs), and coastal development affecting 10,630 reef cells (19.5%). As these pressures have been known for decades, the good news is that a substantial body of conservation science underpins how to design, implement, and measure the success of interventions to achieve fisheries sustainability and improve water quality (Wenger et al. 2017).

While each of the six data layers (**Table 1**) have been demonstrated to alter coral reefs at some magnitude of impact (Darling et al. 2019; Cinner et al. 2016), managing pressures must balance human well-being and ecosystem health, particularly for the millions of people worldwide who rely on coral reefs for livelihoods, culture, food security and coastal protection. Pressures do not need to be ‘removed’ entirely but managed sustainably within complex social, political, and environmental contexts. For example, fisheries management can maintain yields and productivity that delivers a critical source of nutrition and food security to coastal populations (Hicks et al., 2019; McClanahan et al., 2011). Similarly, well-managed coral reef tourism can maintain an estimated US$36 Billion per year global value of coral reef tourism (Spalding et al., 2017). Here, we provide high-resolution spatial maps of local pressures but we highlight that conservation interventions must also address the broader social-economic drivers of the pressures that often operate at larger scales, e.g., multinational investment in fisheries subsidies or global demand for economic development projects in the ‘Blue Economy’ (Bennett et al., 2019).

These results have immediate application for coral reef conservation efforts. Facing the increasing impacts of climate change and most notably severe mass coral bleaching, several major global efforts are taking a ‘refugia-first’ approach, prioritizing conservation and management interventions to potential ‘cool spots’ of climate refuges expected to escape the greatest impacts of bleaching and mass mortality of reef-building corals (e.g., Beyer et al. 2018; Darling et al. 2019). Here, we show that the occurrence of local pressures is similar between refugia and non-refugia locations, providing an opportunity for efforts to mitigate local pressures to scale up beyond climate refugia and to include non-refugia locations as well (e.g., Darling et al., 2019). From local to global scales, these results provide useful information that can be quickly integrated into ongoing coral reef conservation efforts.

Our analysis has three important caveats. First, our analysis only ranks and compares pressures that are available as global data layers. In some locations, important pressures will not be included in our analysis and might require urgent conservation action, such as destructive fishing practices (Bailey & Sumaila, 2015; P. Lestari, pers. comm) or biological invasions by crown of thorn starfish (De’ath, Fabricius, Sweatman, & Puotinen, 2012). This highlights the importance of co-designing conservation interventions with expert knowledge from local resource users and communities, traditional owners, and Indigenous peoples that can effectively incorporate global and local information to prioritize coral reef conservation (Harris et al 2017). Second, our analysis focuses on potential pressures to coral reefs and does not account for ongoing management interventions that are actively managing pressures, such as integrated watershed management or marine protected areas. Strengthening the capacity and resources for management is obviously crucial to ensure pressures do not jeopardize coral reef persistence and ecosystem services and can sustain positive outcomes for biodiversity and coastal communities (Gill et al. 2017). And finally, we do not account for potential interactions among different pressures, which may increase or decrease their realized impacts (Côté, Darling, & Brown, 2016). For example, protection from destructive fishing activities can reduce levels of coral disease, but this benefit is compromised in poor water quality (Lamb et al. 2016).

With the planet’s oceans facing committed warming for decades and predicted to cross a series of climate tipping points (Heinze et al., 2021), ensuring human impacts are effectively managed is urgently required to meet climate adaptation targets of the United Nations Framework Convention on Climate Change and are a key vision of the Post-2020 Biodiversity Framework of the Convention on Biological Diversity. With local pressures expected to expand and intensify over time, particularly with increasing climate change (O’Hara, Frazier, & Halpern 2021), tracking and mitigating these impacts of local pressures also requires access to global data layers (e.g., Halpern 2020). Here, we provide this as underlying datasets, code, and examples of two use cases: pressure report cards and a global mapping platform (**Box 1;** https://programs.wcs.org/vibrantoceans/map). Ensuring the future sustainability of coral reefs requires identifying and managing pressures sustainably with a diverse portfolio of conservation and management interventions. Here, we deliver a comprehensive and highly-resolved analysis of local pressures to coral reefs to guide managers, decision makers and stakeholders towards interventions that ensure this sustainability.

## Acknowledgements and Data

We thank J. Cinner, E. Maire, M. Spalding and L. McLeod for contributing the gravity and tourism value datasets, and the other sources for freely making global data layers available for this study. Funding was provided by Bloomberg Philanthropies Vibrant Oceans Initiative to the Wildlife Conservation Society. Data and code for analysis are publicly accessible at https://github.com/WCS-Marine/local-reef-threats and https://programs.wcs.org/vibrantoceans/map.

## Author Contributions

MA, ED, AW, and GA conceived the study. AW and FS developed data layers. MA and SG conducted the analysis. MA and ED wrote the first draft. All authors edited and approved the manuscript.

## Notes

### Competing Interest Statement

The authors have declared no competing interest.

## References

Allan, J. R., Watson, J. E. M., Marco, M. D., O’Bryan, C. J., Possingham, H. P., Atkinson, S. C., & Venter, O. (2019). Hotspots of human impact on threatened terrestrial vertebrates. PLOS Biology, 17(3), e3000158. doi: 10.1371/journal.pbio.3000158

Bailey, M., & Sumaila, U. R. (2015). Destructive fishing and fisheries enforcement in eastern Indonesia. Marine Ecology Progress Series, 530, 195–211. doi: 10.3354/meps11352

Bennett, N. J., Finkbeiner, E. M., Ban, N. C., Belhabib, D., Jupiter, S. D., Kittinger, J. N., … Christie, P. (2020). The COVID-19 Pandemic, Small-Scale Fisheries and Coastal Fishing Communities. Coastal Management, 48(4), 336–347. doi: 10.1080/08920753.2020.1766937

Bennett, N.J., Cisneros-Montemayor, A.M., Blythe, J., Silver, J.J., Singh, G., Andrews, N., Calò, A., Christie, P., Di Franco, A., Finkbeiner, E.M., & Gelcich, S., 2019. Towards a sustainable and equitable blue economy. Nature Sustainability, 2: 991–993.

Beyer, H. L., Kennedy, E. V., Beger, M., Chen, C. A., Cinner, J. E., Darling, E. S., … Hoegh-Guldberg, O. (2018). Risk-sensitive planning for conserving coral reefs under rapid climate change. Conservation Letters, 11(6), e12587. doi: 10.1111/conl.12587

Borrelli, P., Robinson, D. A., Fleischer, L. R., Lugato, E., Ballabio, C., Alewell, C., … Panagos, P. (2017). An assessment of the global impact of 21st century land use change on soil erosion. Nature Communications, 8(1), 2013. doi: 10.1038/s41467-017-02142-7

Burke, L., Bryant, D., McManus, J., & Spalding, M. (1998). Reefs at Risk. Washington, DC. Retrieved from https://www.wri.org/publication/reefs-risk

Burke, L., Reytar, K., Spalding, M., & Perry, A. (2011). Reefs at Risk Revisited. Washington, DC. Retrieved from https://www.wri.org/publication/reefs-risk-revisited

Center for International Earth Science Information Network - CIESIN - Columbia University. (2018). Gridded Population of the World, Version 4 (GPWv4): Population Count Adjusted to Match 2015 Revision of UN WPP Country Totals, Revision 11. Palisades, NY: NASA Socioeconomic Data and Applications Center (SEDAC). Retrieved from https://doi.org/10.7927/H4PN93PB

Cinner, J. E., Huchery, C., MacNeil, M. A., Graham, N. A. J., McClanahan, T. R., Maina, J., … Mouillot, D. (2016). Bright spots among the world’s coral reefs. Nature, 535(7612), 416–419. doi: 10.1038/nature18607

Cinner, J. E., Maire, E., Huchery, C., MacNeil, M. A., Graham, N. A. J., Mora, C., … Mouillot, D. (2018). Gravity of human impacts mediates coral reef conservation gains. Proceedings of the National Academy of Sciences, 201708001. doi: 10.1073/pnas.1708001115

Claar, D. C., Starko, S., Tietjen, K. L., Epstein, H. E., Cunning, R., Cobb, K. M., … Baum, J. K. (2020). Dynamic symbioses reveal pathways to coral survival through prolonged heatwaves. Nature Communications, 11(1), 6097. doi: 10.1038/s41467-020-19169-y

Côté, I. M., Darling, E. S., & Brown, C. J. (2016). Interactions among ecosystem stressors and their importance in conservation. Proceedings of the Royal Society B: Biological Sciences, 283(1824), 20152592. doi: 10.1098/rspb.2015.2592

Darling, E. S., McClanahan, T. R., Maina, J., Gurney, G. G., Graham, N. A. J., Januchowski-Hartley, F., … Mouillot, D. (2019). Social–environmental drivers inform strategic management of coral reefs in the Anthropocene. Nature Ecology & Evolution, 3(9), 1341–1350. doi: 10.1038/s41559-019-0953-8

De’ath, G., Fabricius, K. E., Sweatman, H., & Puotinen, M. (2012). The 27-year decline of coral cover on the Great Barrier Reef and its causes. Proceedings of the National Academy of Sciences of the United States of America, 109(44), 17995–17999. doi: 10.1073/pnas.1208909109

Eddy, T. D., Cheung, W. W. L., & Bruno, J. F. (2018). Historical baselines of coral cover on tropical reefs as estimated by expert opinion. PeerJ, 6, e4308. doi: 10.7717/peerj.4308

Grantham, H. S., Duncan, A., Evans, T. D., Jones, K. R., Beyer, H. L., Schuster, R., … Watson, J. E. M. (2020). Anthropogenic modification of forests means only 40% of remaining forests have high ecosystem integrity. Nature Communications, 11(1), 5978. doi: 10.1038/s41467-020-19493-3

Halpern, B. S. (2020). Building on a Decade of the Ocean Health Index. One Earth, 2(1), 30–33. doi: 10.1016/j.oneear.2019.12.011

Harris, J. L., Estradivari, E., Fox, H. E., McCarthy, O. S., & Ahmadia, G. N. (2017). Planning for the future: Incorporating global and local data to prioritize coral reef conservation. Aquatic Conservation: Marine and Freshwater Ecosystems, 27(S1), 65–77. doi: https://doi.org/10.1002/aqc.2810

Heinze, C., Blenckner, T., Martins, H., Rusiecka, D., Döscher, R., Gehlen, M., … Wilson, S. (2021). The quiet crossing of ocean tipping points. Proceedings of the National Academy of Sciences, 118(9). doi: 10.1073/pnas.2008478118

Hicks, C. C., Cohen, P. J., Graham, N. A. J., Nash, K. L., Allison, E. H., D’Lima, C., … MacNeil, M. A. (2019). Harnessing global fisheries to tackle micronutrient deficiencies. Nature, 574(7776), 95–98. doi: 10.1038/s41586-019-1592-6

Hoegh-Guldberg, O., Jacob, D., Taylor, M., Bindi, M., Brown, S., Camilloni, I., … Sherstyukov, B. (2018). Impacts of 1.5°C of Global Warming on Natural and Human Systems. In V. Masson-Delmotte, P. Zhai, & H.O. Pörtner, Global Warming of 1.5°C. An IPCC Special Report on the impacts of global warming of 1.5°C above pre-industrial levels and related global greenhouse gas emission pathways, in the context of strengthening the global response to the threat of climate change, sustainable development, and efforts to eradicate poverty (p. 138). Intergovernmental Panel on Climate Change.

Hughes, T. P., Kerry, J. T., Baird, A. H., Connolly, S. R., Dietzel, A., Eakin, C. M., … Torda, G. (2018). Global warming transforms coral reef assemblages. Nature, 556(7702), 492–496. doi: 10.1038/s41586-018-0041-2

King, C., Iba, W., & Clifton, J. (2021). Reimagining resilience: COVID-19 and marine tourism in Indonesia. Current Issues in Tourism, 0(0), 1–17. doi: 10.1080/13683500.2021.1873920

Lamb, J. B., Wenger, A. S., Devlin, M. J., Ceccarelli, D. M., Williamson, D. H., & Willis, B. L. (2016). Reserves as tools for alleviating impacts of marine disease. Philosophical Transactions of the Royal Society B: Biological Sciences, 371(1689), 20150210. doi: 10.1098/rstb.2015.0210

Maina, J., de Moel, H., Zinke, J., Madin, J., McClanahan, T., & Vermaat, J. E. (2013). Human deforestation outweighs future climate change impacts of sedimentation on coral reefs. Nature Communications, 4, 1986. doi: 10.1038/ncomms2986

McClanahan, T. R., Graham, N. A. J., MacNeil, M. A., Muthiga, N. A., Cinner, J. E., Bruggemann, J. H., & Wilson, S. K. (2011). Critical thresholds and tangible targets for ecosystem-based management of coral reef fisheries. Proceedings of the National Academy of Sciences, 108(41), 17230–17233. doi: 10.1073/pnas.1106861108

Morrison, T. H., Hughes, T. P., Adger, W. N., Brown, K., Barnett, J., & Lemos, M. C. (2019). Save reefs to rescue all ecosystems. Nature, 573(7774), 333–336. doi: 10.1038/d41586-019-02737-8

O’Hara, C. C., Frazier, M., & B. S. Halpern. (2021). At-risk marine biodiversity faces extensive, expanding, and intensifying human impacts. Science, 372: 84–87.

R Core Team. (2019). R: A Language and Environment for Statistical Computing. R Foundation for Statistical Computing, Vienna, Austria. Retrieved from http://www.R-project.org/

Spalding, M., Burke, L., Wood, S. A., Ashpole, J., Hutchison, J., & zu Ermgassen, P. (2017). Mapping the global value and distribution of coral reef tourism. Marine Policy, 82, 104–113. doi: 10.1016/j.marpol.2017.05.014

Tulloch, V. J., Tulloch, A. I., Visconti, P., Halpern, B. S., Watson, J. E., Evans, M. C., … Possingham, H. P. (2015). Why do we map threats? Linking threat mapping with actions to make better conservation decisions. Frontiers in Ecology and the Environment, 13(2), 91–99. doi: 10.1890/140022

Tulloch, V. J., Turschwell, M. P., Giffin, A. L., Halpern, B. S., Connolly, R., Griffiths, L., … Brown, C. J. (2020). Linking threat maps with management to guide conservation investment. Biological Conservation, 245, 108527. doi: 10.1016/j.biocon.2020.108527

UNEP-WCMC, WorldFish Centre, W.I. & TNC. (2010). Global distribution of warm-water coral reefs, compiled from multiple sources including the Millennium Coral Reef Mapping Project. Version 1.3. Retrieved from http://data.unep-wcmc.org/datasets/1

Wear, S. L. (2016). Missing the boat: Critical threats to coral reefs are neglected at global scale. Marine Policy, 74, 153–157. doi: 10.1016/j.marpol.2016.09.009

Wear, S. L., & Thurber, R. V. (2015). Sewage pollution: Mitigation is key for coral reef stewardship. Annals of the New York Academy of Sciences, 1355(1), 15–30. doi: 10.1111/nyas.12785

Wenger, A. S., Harris, D., Weber, S., Vaghi, F., Nand, Y., Naisilisili, W., … Jupiter, S. D. (2020). Best-practice forestry management delivers diminishing returns for coral reefs with increased land-clearing. Journal of Applied Ecology, 57(12), 2381–2392. doi: https://doi.org/10.1111/1365-2664.13743

Wenger, A. S., Harvey, E., Wilson, S., Rawson, C., Newman, S. J., Clarke, D., … Evans, R. D. (2017). A critical analysis of the direct effects of dredging on fish. Fish and Fisheries, 18(5), 967–985. doi: https://doi.org/10.1111/faf.12218

Wenger, A. S., Rawson, C. A., Wilson, S., Newman, S. J., Travers, M. J., Atkinson, S., … Harvey, E. (2018). Management strategies to minimize the dredging impacts of coastal development on fish and fisheries. Conservation Letters, 11(5), e12572. doi: 10.1111/conl.12572

Yap, W. Y., & Lam, J. S. L. (2013). 80 million-twenty-foot-equivalent-unit container port? Sustainability issues in port and coastal development. Ocean & Coastal Management, 71, 13–25. doi: 10.1016/j.ocecoaman.2012.10.011

