## Supporting Information for "A global map of human pressures on tropical coral reefs"

### Supplementary Methods

#### Definitions

**Bioclimatic unit, BCU:** A reef area of ~500 km$^2$ identified as part of a global portfolio for potential global climate refugia for coral reefs (Beyer et al., 2018). BCUs were identified to maximise a set of climatic criteria, comprising: historical thermal stress, recent thermal stress, future thermal stress, cyclone impacts and larval connectivity. On average, there are 220 coral reef cells per BCU (median; interquartile range: 111 – 311).

**Pressure:** A contextual variable that can, in certain circumstances, exert unsustainable impact on a reef and degrade its ecological function, productivity, or resilience. While each variable could result in negative consequences for reefs, they are not necessarily activities that need to be stopped, but rather managed to ensure sustainability over the long term. For example, fishing is important for food and livelihood security but at high levels of exploitation can shift coral reefs to low fish biomass and often algae-dominated reefs. Similarly, tourism can cause direct damage to corals from people stepping on the reef, more pollution from coastal development of hotels, piers, etc. with limited waste management, or more fisheries exploitation from sport fishing by tourists or increased local fishing to meet tourist demand for fresh seafood. However, tourism can also be an important factor enabling conservation, e.g., by supporting alternative livelihoods for a positive conservation impact. All variables should be considered appropriately in each context, and refined with local knowledge.

#### Pressure data layers

**Fishing.** Pressures due to fishing were measured as market gravity. The gravity concept draws on an analogy from Newton's Law of Gravitation and predicts that interactions between two places (e.g., cities) are positively related to their mass (i.e., population) and inversely related to the distance between them (Anderson, 2011). Cinner et al. (2018) applied the gravity concept to coral reefs under the assumption that human interactions with a reef are a function of the population of a place divided by the squared time it takes to travel to the reefs. Using travel time instead of linear distance accounts for the differences incurred by traveling over different surfaces, such as water, roads or tracks (Maire et al., 2016). High market gravity, measured as (number of people) / (hours of travel)$^2$, reduces fish biomass and the occurrence of top predators on coral reefs. As noted by Cinner et al. (2018), gravity values can be similar for places that have large populations but are far from the reefs (e.g., population = 15,000 people, travel time = 7 h, gravity = 306) as to those with small populations that are close to the reef (e.g., population = 300 people, travel time = 1 h, gravity = 300. We mapped gravity to the coral reef grid cells by calculating the mean value of market gravity of all the intersecting grid cells from Cinner et al.'s original layer (see R code).

**Coastal development.** Coastal development has profound effects on nearshore ecosystems through wastewater discharge (Wear & Thurber, 2015), construction (Wenger et al., 2017), aquaculture, industry and agriculture. As an indicator of coastal development, we used the number of people living on the coast obtained from the population count layer v 4.11 (Center for International Earth Science Information Network - CIESIN - Columbia University, 2018) for the year 2020 at a spatial resolution of 2.5 minutes of degree (about 5 km). Population counts were mapped to the grid cells by summing their values within a distance of 5 km from the boundaries of each grid cell. We included population counts within 5 km distance on the basis of studies that have recorded recovery of coral reefs following wastewater management (Birkeland, Green, Fenner, Squair, & Dahl, 2013; Hunter & Evans, 1995). Using different radii (1 km to 50 km) resulted in strongly correlated values (Spearman's correlation coefficient > 0.7) for this pressure (**Figure S12**).

**Industrial development.** Pressures from industrial development are a special type of coastal development pressures associated with maritime activities such as dredging. We used location of ports as a proxy for dredging, based on the prevalence of dredging activities at port facilities (Yap & Lam, 2013). We sourced port locations, in the form of point data, from a spatial layer of all existing ports, freely available from Google data (https://goo.gl/Yu8xxt). We excluded all inland ports and any duplicates, resulting in 2,646 coastal ports. Values indicate the number of ports within 5km from the boundary of each coral reef grid cell. We selected ports within 5 km as this distance reflected the maximum likely spatial extent of dredging impacts, acknowledging that the spatial extent of any dredging operation will be dependent on local conditions, including dredge type, material disposal, and local currents (Wenger et al., 2018).

**Tourism.** Coral reef related tourism is an important industry providing employment and income, but is not always benign. Negative impacts can include degradation and loss of marine life through diving and snorkeling; indirect impacts arising from poorly planned coastal development (construction, sewage, solid waste and dredging in remote locations); and increasing levels of coastal demand for seafood and curios (Albuquerque et al., 2014). Spalding et al. (2017) estimated the monetary value of domestic and international tourism on coral reefs accounting for both direct non-extractive uses and activities indirectly linked to the presence of nearby reefs (e.g. the role of reefs in generating clear calm waters and beach sand, outstanding views, etc.). We mapped reef value to the coral reef grid cells by summing the reef values of all the intersecting grid cells from Spalding et al.'s original layer. Reef value is expressed in 1000 US$ per square km of reef.

**Pollution: sedimentation**. We combined two approaches for estimating sediment delivery to river mouths. For the first approach, we estimated sediment delivery using the Natural Capital Project's Integrated Valuation of Ecosystem Service Sediment Delivery Ratio (SDR) Model (InVEST SDR version 3.7.0) (Hamel, Chaplin-Kramer, Sim, & Mueller, 2015; Tallis & Polasky, 2009). Operating at the resolution of the land-cover input (250m), the SDR model calculates sediment yield by combining the revised universal soil loss equation (RUSLE) with a sediment delivery ratio (SDR) to quantify the amount of soil eroded for a given area that will travel to a stream pour-point (Hamel et al., 2015). The RUSLE approach uses five different environmental input factors: land cover and management factor (C-factor), rainfall-runoff erosivity (R-factor), slope length and steepness factor (LS-factor), soil erodibility (K-factor). Each of these layers plus global land cover data were taken from Borrelli et al. (2017). For the second approach, we used the SDR values from the Reefs at Risk analysis (Burke, Reytar, Spalding, & Perry, 2011). While the calculations for these SDR values are more simplistic than the InVEST model, they account for sediment trapping potential of dams and mangroves, which InVEST does not.

The sediment exposure to adjacent reefs was modelled using a sediment plume model, written and run in R version 3.5.3. The dispersal of potential sediment at each river mouth was modelled using a cost-path surface, where a decay function evenly distributes 0.5% of the initial potential sediment value to all adjacent cells, until either a threshold of 0.05% of the global maximum value or a distance of 80 km from the river mouth was reached (Burke et al., 2011; Halpern et al., 2008). The sediment plumes from each river mouth were then summed to provide a cumulative sediment exposure value, measured as tons of sediment / $km^2$. Finally, the sediment plumes for the two approaches were averaged.

**Pollution: nitrogen.** Nitrogen delivery to the river mouth was calculated using a modified method developed from Halpern et al. (2008). The proportional crop cover in each reef catchment (*n* = 97,255) was compared to the country crop cover, using the land cover layers in Borrelli et al. (2017). The total annual nitrogen fertilizer use for each country reported by the Food and Agriculture Organization (FAO) was distributed to each reef catchment, based on the proportional crop cover. Tropical plant uptake of fertilizer N has been estimated to be 50% of the applied quantity, but can vary from between 20 to 80% (Baligar & Bennett, 1986). Although wastewater discharged into riverine environments will also lead to nitrogen pollution runoff, we were not able to account for this source of nitrogen. We estimated nitrogen delivery to the river mouth for both a 20% uptake and an 80% uptake of nitrogen fertilizer.

The nitrogen plume was modeled using a modified version of the sediment plume dispersal model. We dispersed the nitrogen at the river mouth using both the decay function used by the Ocean Health Index (0.5%) and a decay function of 0.25% to account for observed conservative mixing and greater transport of nutrients in the marine environment (Devlin et al., 2013). The nitrogen plumes from each river mouth were then summed to provide a cumulative nitrogen exposure value. The plumes for each scenario (the two uptake rate scenarios and the two plume decay function scenarios) were then averaged.

**Cumulative impact score.** A cumulative pressure impact score for each reef pixel was estimated from the weighted average of the percentiles of the six pressure layers. Weights were chosen from Wear (2016) who conducted a survey of 170 managers from 50 coral reef countries and territories to calculate the perceived severity of different pressures on coral reefs (see **Table 1** in the main text). We also calculated the cumulative impact score from all pressures weighted equally (e.g., a mean average pressure percentile) and this was highly correlated to the weighted index (Spearman's correlation coefficient $r$ = 0.99, p-value < 2.2e-16); we thus use the weighted cumulative impact score for further analyses.

#### Ranking top pressures using percentiles

After summarizing the value of each pressure within 5km x 5km reef pixels, we calculate the percentile of each pressure within each pixel, relative to the global distribution. Here, a pixel with a 0% percentile would be the lowest value in the data layer, and a pixel with a 100% percentile has the highest value. Using percentiles allowed us to standardize and compare across pressures within each reef pixel. The top-ranked pressure was identified as the data layer with the highest percentile.

#### Coding

We performed all mapping and data analyses in R 3.6.0 (R Core Team, 2019) using packages 'ggpubr' 0.4.0, 'ggbeeswarm' 0.6.0, 'magick' 2.6.0, 'rmarkdown' 2.6, 'sf' 0.9-6, 'tidyverse' 1.3.0 and 'tmap' 2.3-1 and tmaptools 3.1 (Clarke & Sherrill-Mix, 2017; Kassambara, 2020; Ooms, 2021; Pebesma, 2018; Tennekes, 2018, 2021; Wickham et al., 2019; Xie, Allaire, & Grolemund, 2020). The R code and underlying data layers are available at https://github.com/WCS-Marine/local-reef-threats.

### References

Albuquerque, T., Loiola, M., Nunes, J. de A. C. C., Reis-Filho, J. A., Sampaio, C. L. S., & Leduc, A. O. H. C. (2014). In situ effects of human disturbances on coral reef-fish assemblage structure: Temporary and persisting changes are reflected as a result of intensive tourism. *Marine and Freshwater Research*, *66*(1), 23–32. doi: 10.1071/MF13185

Anderson, J. E. (2011). The Gravity Model. *Annual Review of Economics*, *3*(1), 133–160. doi:

10.1146/annurev-economics-111809-125114

Baligar, V., & Bennett, O. (1986). NPK-fertilizer efficiency—A situation analysis for the tropics. *Fertilizer Research*, *10*(2), 147–164. doi: 10.1007/BF01074369

Beyer, H. L., Kennedy, E. V., Beger, M., Chen, C. A., Cinner, J. E., Darling, E. S., … Hoegh-Guldberg, O. (2018). Risk-sensitive planning for conserving coral reefs under rapid climate change. *Conservation Letters*, *11*(6), e12587. doi: 10.1111/conl.12587

Birkeland, C., Green, A., Fenner, D., Squair, C., & Dahl, A. L. (2013). Substratum stability and coral reef resilience: Insights from 90 years of disturbances on a reef in American Samoa. *Micronesica*, (6), 1–16.

Borrelli, P., Robinson, D. A., Fleischer, L. R., Lugato, E., Ballabio, C., Alewell, C., … Panagos, P. (2017). An assessment of the global impact of 21st century land use change on soil erosion. *Nature Communications*, *8*(1), 2013. doi: 10.1038/s41467-017-02142-7

Burke, L., Reytar, K., Spalding, M., & Perry, A. (2011). *Reefs at Risk Revisited*. Washington, DC. Retrieved from https://www.wri.org/publication/reefs-risk-revisited

Center for International Earth Science Information Network - CIESIN - Columbia University. (2018). *Gridded Population of the World, Version 4 (GPWv4): Population Count Adjusted to Match 2015 Revision of UN WPP Country Totals, Revision 11*. Palisades, NY: NASA Socioeconomic Data and Applications Center (SEDAC). Retrieved from https://doi.org/10.7927/H4PN93PB

Cinner, J. E., Maire, E., Huchery, C., MacNeil, M. A., Graham, N. A. J., Mora, C., … Mouillot, D. (2018). Gravity of human impacts mediates coral reef conservation gains. *Proceedings of the National Academy of Sciences*, 201708001. doi: 10.1073/pnas.1708001115

Clarke, E., & Sherrill-Mix, S. (2017). ggbeeswarm: Categorical Scatter (Violin Point) Plots (Version 0.6.0). Retrieved from https://CRAN.R-project.org/package=ggbeeswarm

Devlin, M. J., da Silva, E. T., Petus, C., Wenger, A., Zeh, D., Tracey, D., … Brodie, J. (2013). Combining in-situ water quality and remotely sensed data across spatial and temporal scales to measure variability in wet season chlorophyll-a: Great Barrier Reef lagoon (Queensland, Australia). *Ecological Processes*, *2*(1), 31. doi: 10.1186/2192-1709-2-31

Halpern, B. S., Walbridge, S., Selkoe, K. A., Kappel, C. V., Micheli, F., D'Agrosa, C., … Watson, R. (2008). A global map of human impact on marine ecosystems. *Science*, *319*(5865), 948–952. (WOS:000253165700045). doi: 10.1126/science.1149345

Hamel, P., Chaplin-Kramer, R., Sim, S., & Mueller, C. (2015). A new approach to modeling the sediment retention service (InVEST 3.0): Case study of the Cape Fear catchment, North Carolina, USA. *Science of The Total Environment*, *524–525*, 166–177. doi: 10.1016/j.scitotenv.2015.04.027

Hunter, C. L., & Evans, C. W. (1995). Coral Reefs in Kaneohe Bay, Hawaii: Two Centuries of Western Influence and Two Decades of Data. *Bulletin of Marine Science*, *57*(2), 501–515.

Kassambara, A. (2020). ggpubr: "ggplot2" Based Publication Ready Plots (Version 0.4.0). Retrieved from https://CRAN.R-project.org/package=ggpubr

Maire, E., Cinner, J., Velez, L., Huchery, C., Mora, C., Dagata, S., … Mouillot, D. (2016). How accessible are coral reefs to people? A global assessment based on travel time. *Ecology Letters*, *19*(4), 351–360. doi: 10.1111/ele.12577

Ooms, J. (2021). magick: Advanced Graphics and Image-Processing in R (Version 2.7.1). Retrieved from https://CRAN.R-project.org/package=magick

Pebesma, E. (2018). Simple Features for R: Standardized Support for Spatial Vector Data. *The R Journal*, *10*(1), 439–446.

Spalding, M., Burke, L., Wood, S. A., Ashpole, J., Hutchison, J., & zu Ermgassen, P. (2017). Mapping the global value and distribution of coral reef tourism. *Marine Policy*, *82*, 104–113. doi: 10.1016/j.marpol.2017.05.014

Tallis, H., & Polasky, S. (2009). Mapping and valuing ecosystem services as an approach for conservation and natural-resource management. *Annals of the New York Academy of Sciences*, *1162*, 265–283. doi: 10.1111/j.1749-6632.2009.04152.x

Tennekes, M. (2018). tmap: Thematic Maps in R. *Journal of Statistical Software*, *84*(1), 1–39. doi: 10.18637/jss.v084.i06

Tennekes, M. (2021). tmaptools: Thematic Map Tools (Version 3.1-1). Retrieved from https://CRAN.R-project.org/package=tmaptools

Wear, S. L., & Thurber, R. V. (2015). Sewage pollution: Mitigation is key for coral reef stewardship. *Annals of the New York Academy of Sciences*, *1355*(1), 15–30. doi: 10.1111/nyas.12785

Wenger, A. S., Harvey, E., Wilson, S., Rawson, C., Newman, S. J., Clarke, D., … Evans, R. D. (2017). A critical analysis of the direct effects of dredging on fish. *Fish and Fisheries*, *18*(5), 967–985. doi: https://doi.org/10.1111/faf.12218

Wenger, A. S., Rawson, C. A., Wilson, S., Newman, S. J., Travers, M. J., Atkinson, S., … Harvey, E. (2018). Management strategies to minimize the dredging impacts of coastal development on fish and fisheries. *Conservation Letters*, *11*(5), e12572. doi: 10.1111/conl.12572

Wickham, H., Averick, M., Bryan, J., Chang, W., McGowan, L. D., François, R., … Yutani, H. (2019). Welcome to the Tidyverse. *Journal of Open Source Software*, *4*(43), 1686. doi: 10.21105/joss.01686

Xie, Y., Allaire, J. J., & Grolemund, G. (2020). *R Markdown: The Definitive Guide*. Retrieved from https://bookdown.org/yihui/rmarkdown/

Yap, W. Y., & Lam, J. S. L. (2013). 80 million-twenty-foot-equivalent-unit container port? Sustainability issues in port and coastal development. *Ocean & Coastal Management*, *71*, 13–25. doi: 10.1016/j.ocecoaman.2012.10.011

#### Box 1

We prepared pressure report cards for each BCU, publicly available at https://github.com/WCS-Marine/local-reef-threats. Each report card includes a regional map showing the location of the BCU with major geographical elements (cities, names of marine regions, islands, etc.) and a set of detailed maps of the six different pressures showing the variability of the pressure across the specified BCU. Within-BCU variability is also shown in a beeswarm plot (see Figure): each dot shows the value for one reef cell, and together they show the variability of the values within the BCU. The colour of the dots is proportional to the pressure percentile. In case of high within-BCU variability (as for tourism in this example), the median could be zero while a few reef cells can have higher percentiles. Note that reef cells with zero values are all tied to a zero percentile (blue dots).

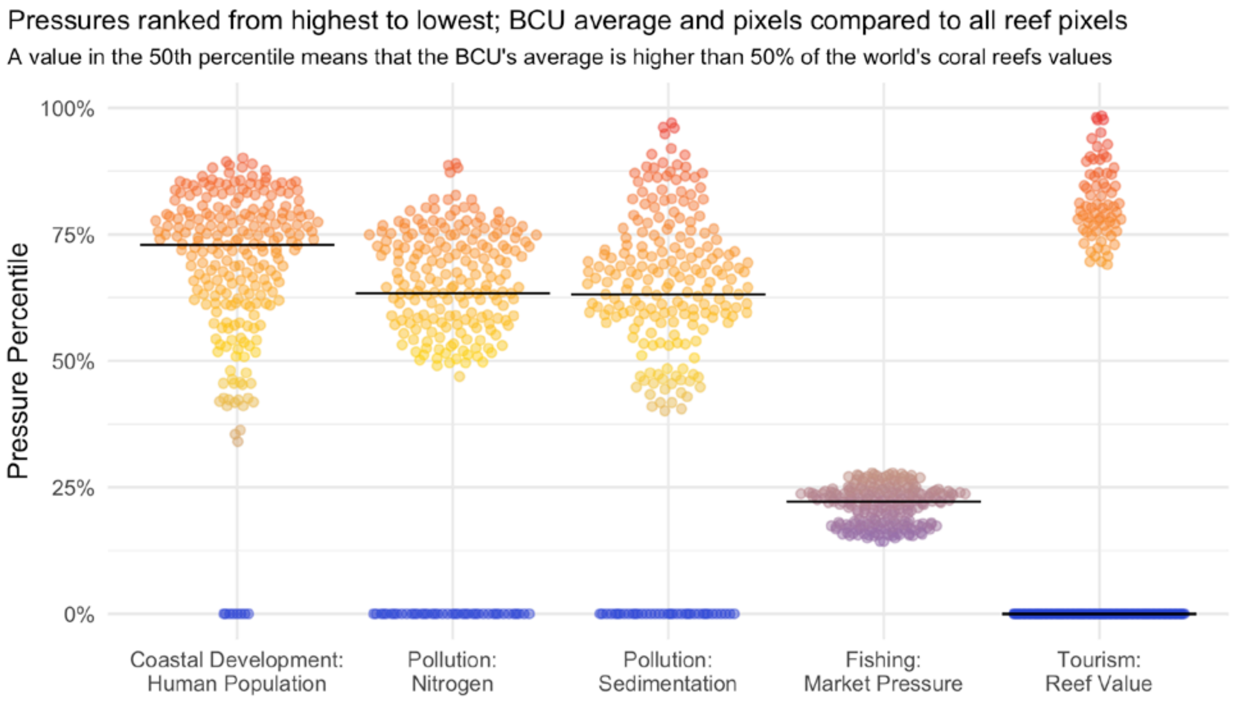


[End of Box 1]

#### Supplementary Figures

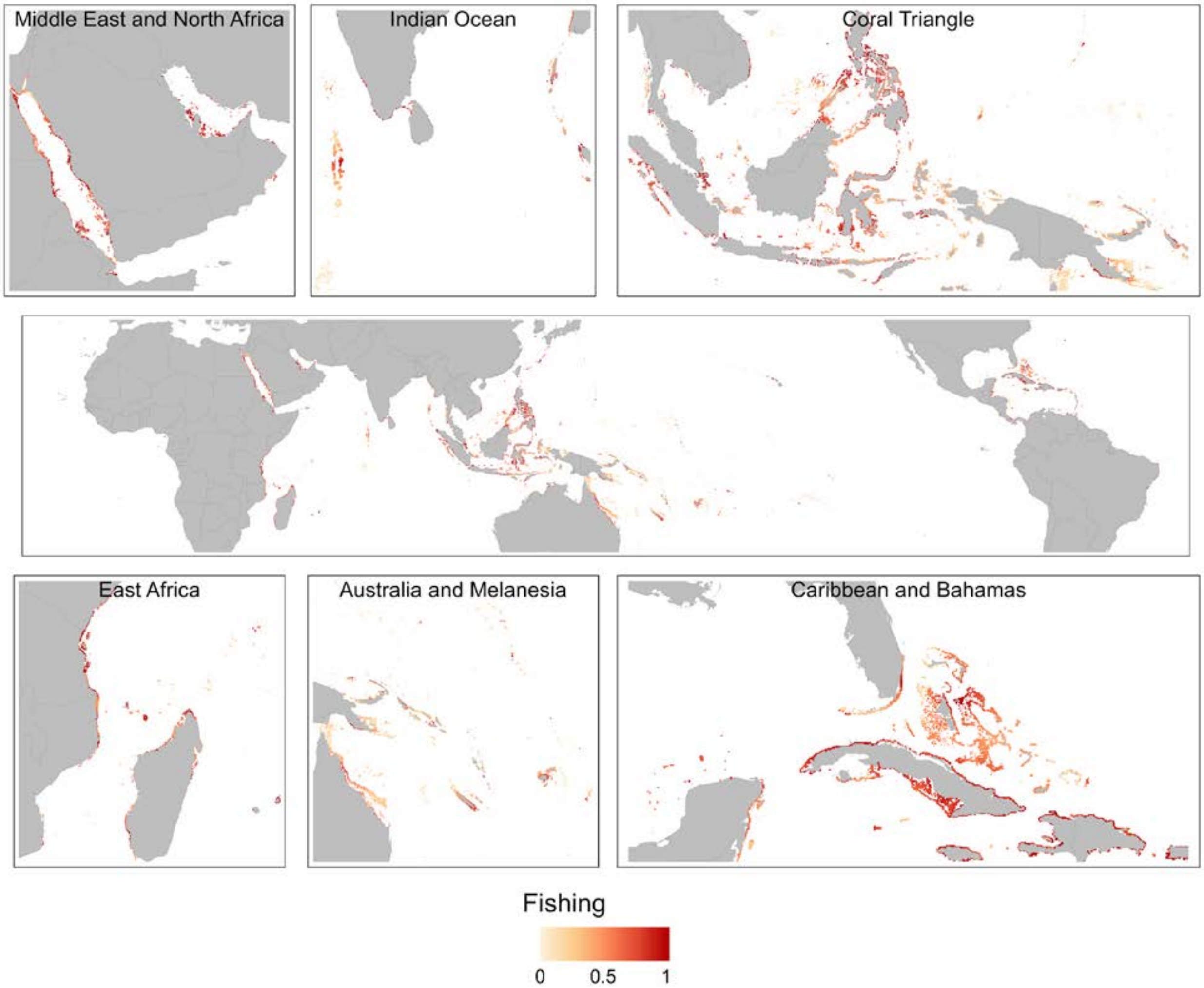


**Figure S1.** Global map of fishing pressure data layer, with insets by key coral reef regions. Values are distributed from 0 (lowest value in data layer) to 1 (highest value in data layer).

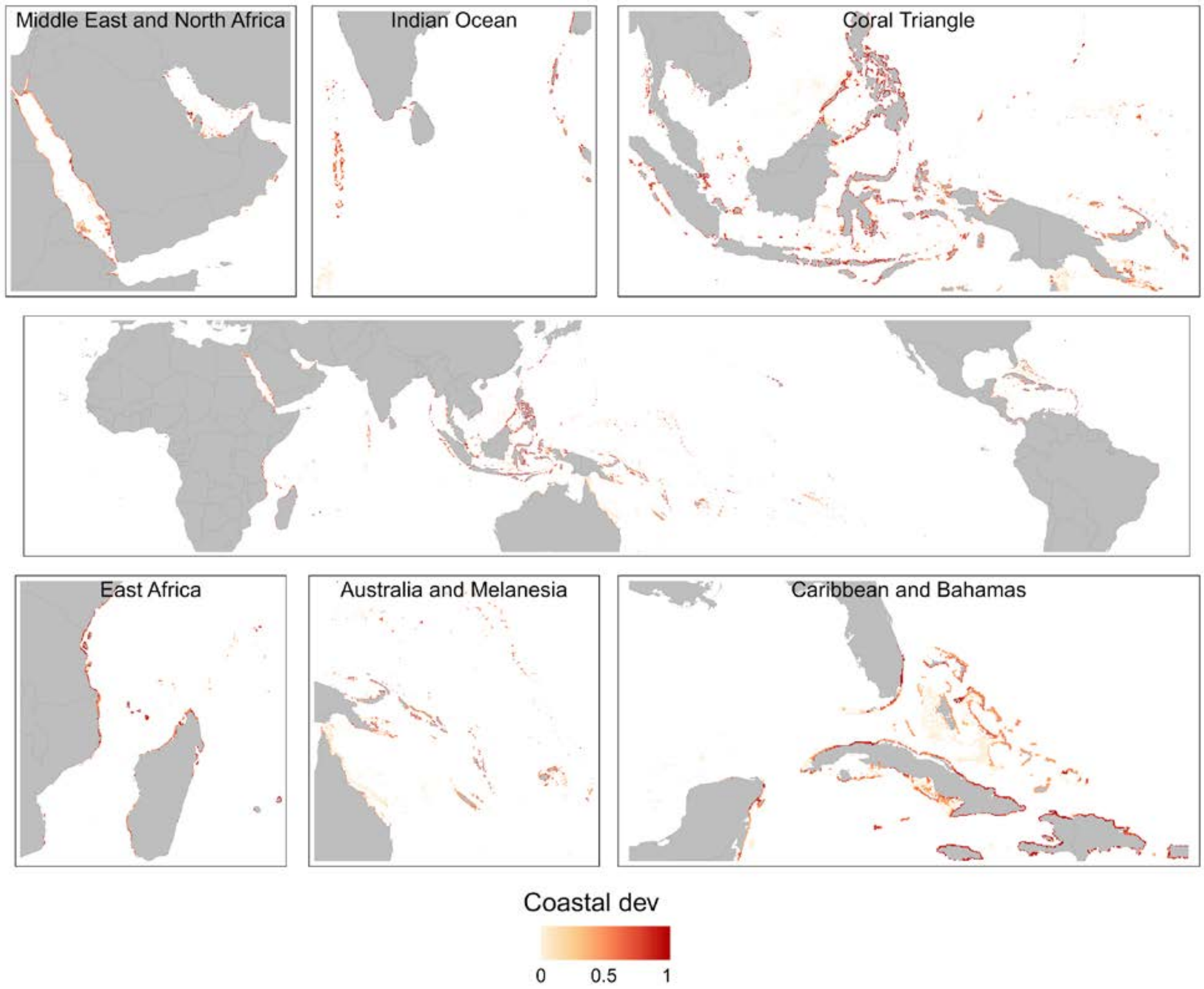


**Figure S2.** Global map of coastal development (human population) data layer, with insets by key coral reef regions. Values are distributed from 0 (lowest value in data layer) to 1 (highest value in data layer).

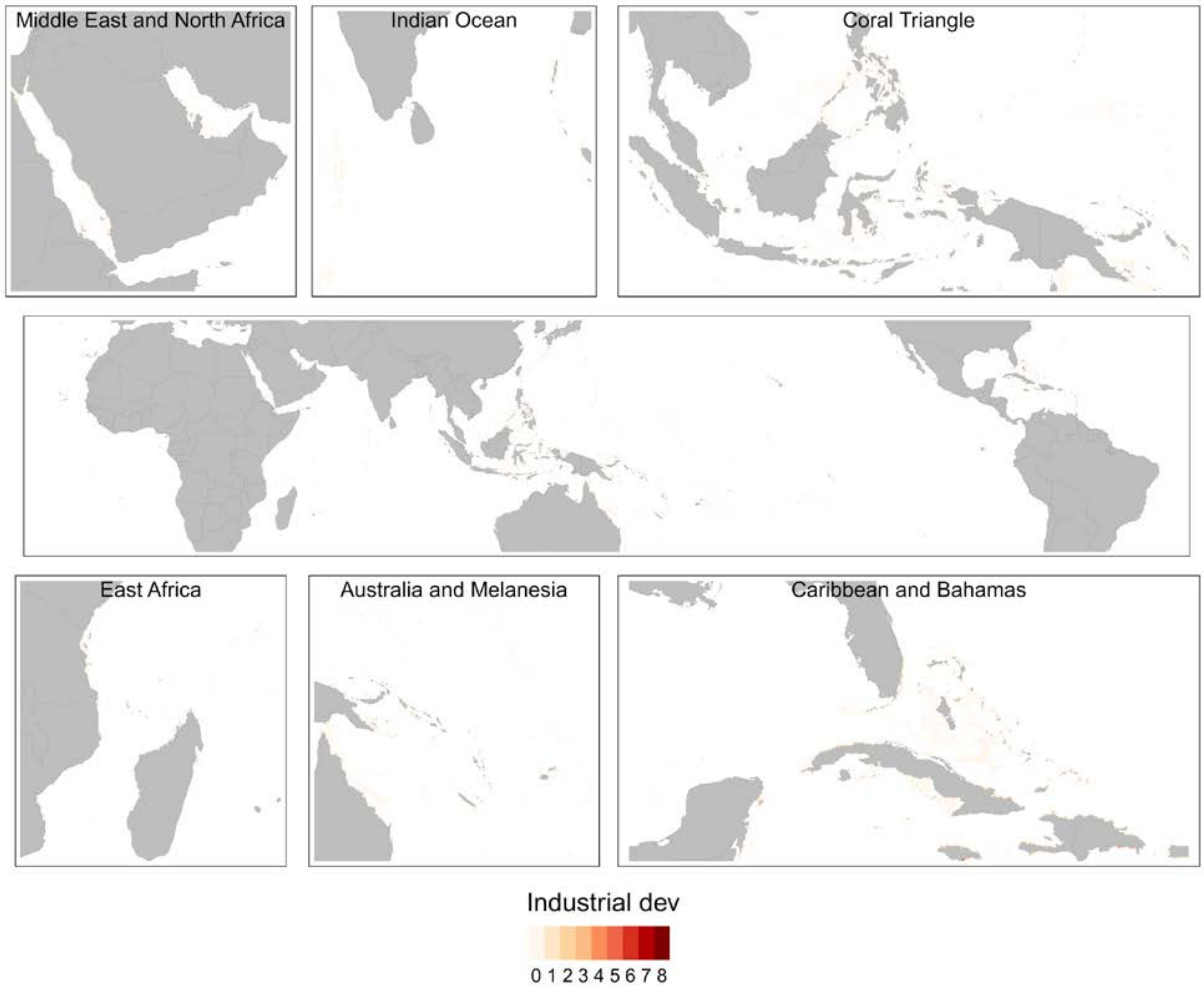


**Figure S3.** Global map of industrial development (number of ports) data layer, with insets by key coral reef regions. Values are the number of industrial ports in a reef cell.

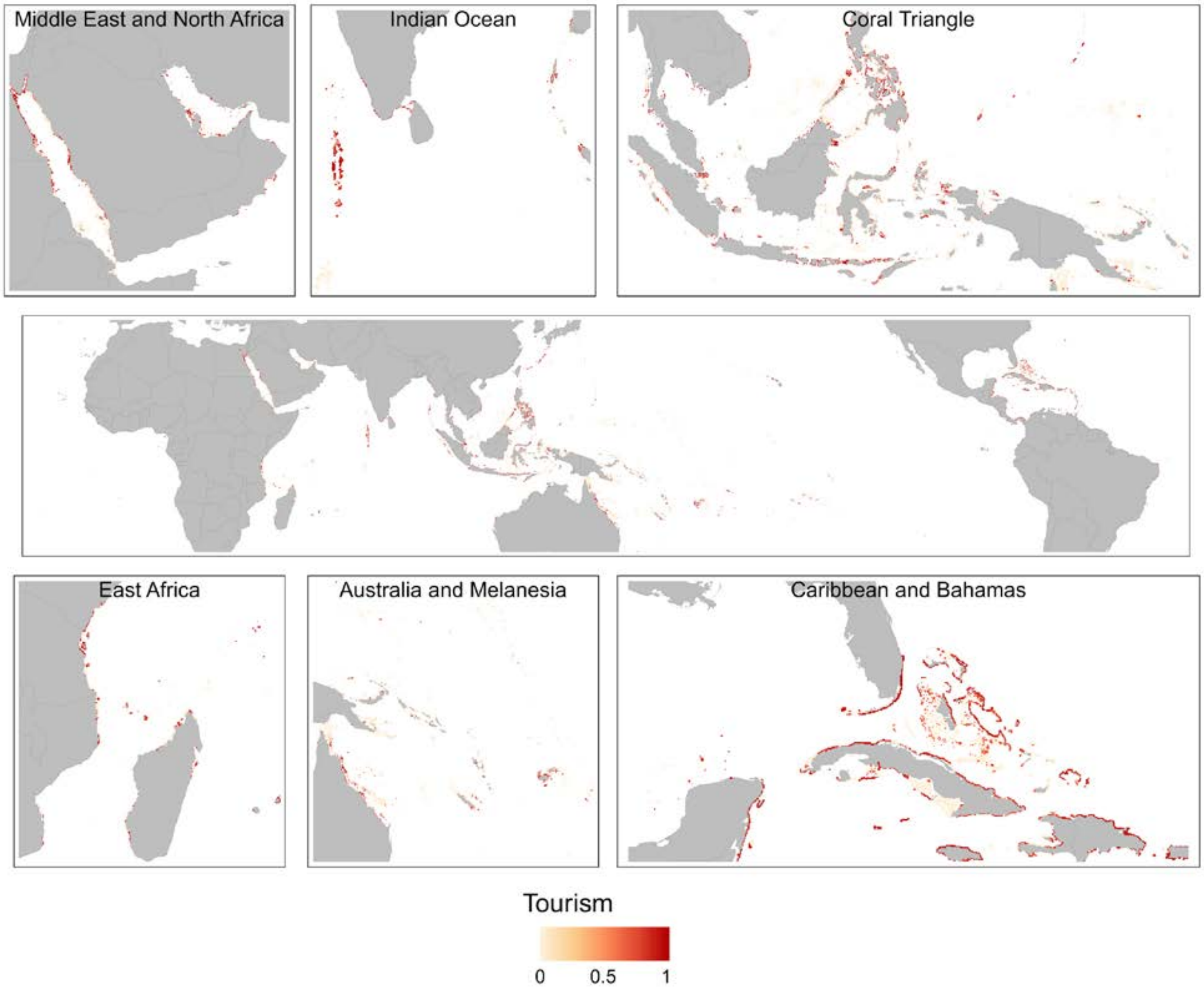


**Figure S4.** Global map of coral reef tourism values ($/km$^2$) data layer, with insets by key coral reef regions. Values are distributed from 0 (lowest value in data layer) to 1 (highest value in data layer).

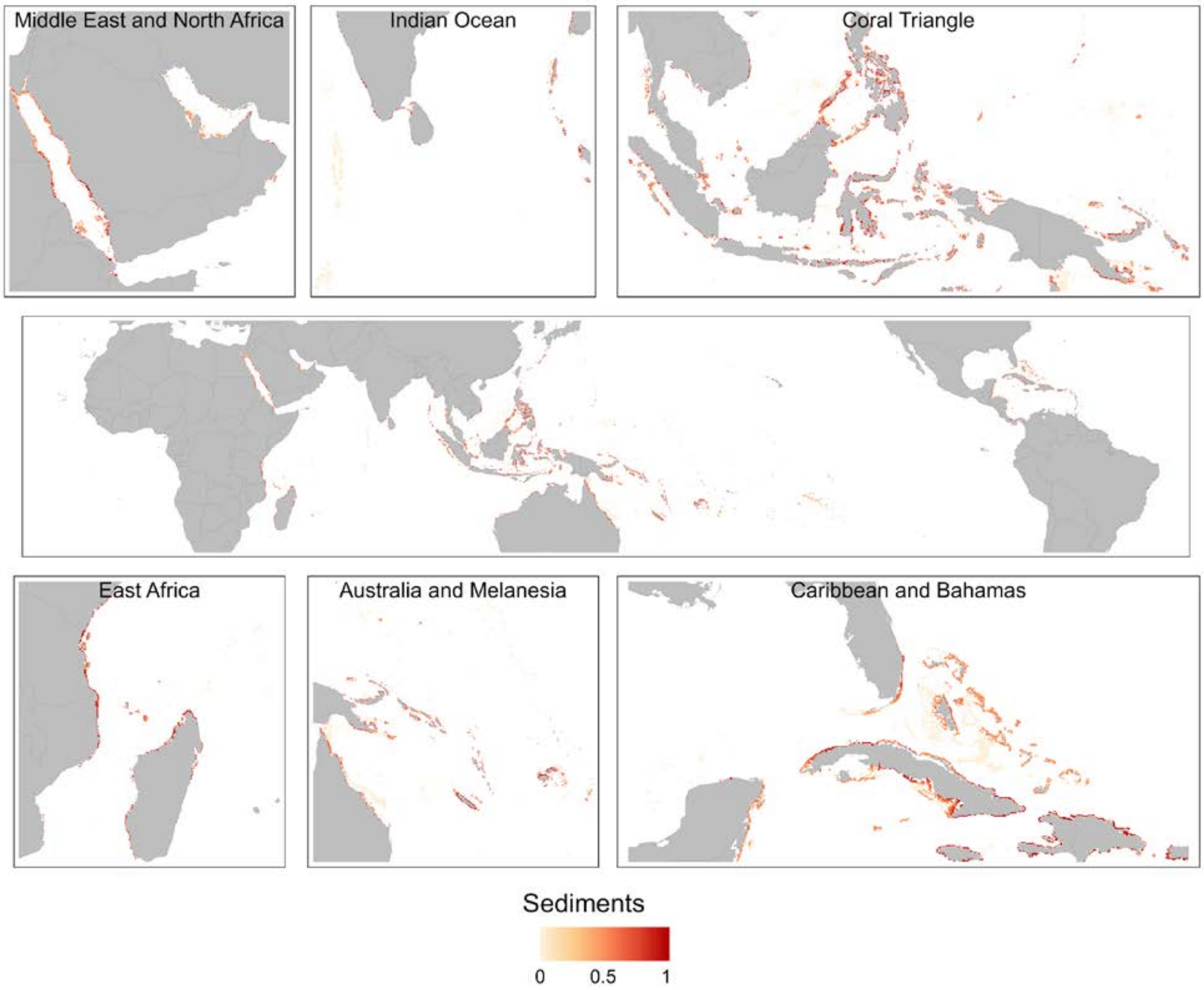


**Figure S5.** Global map of water pollution: sediments data layer, with insets by key coral reef regions. Values are distributed from 0 (lowest value in data layer) to 1 (highest value in data layer).

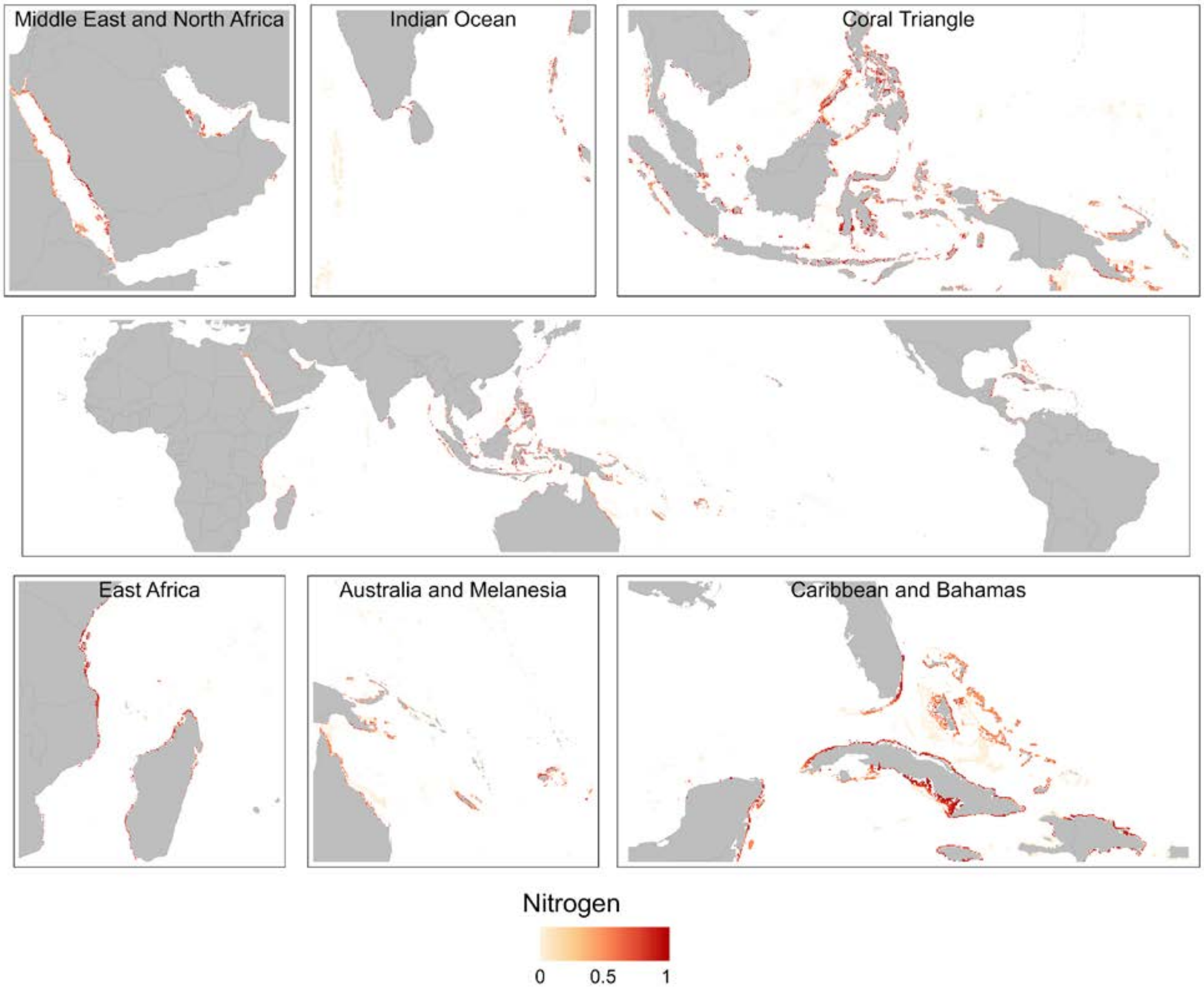


**Figure S6.** Global map of water pollution: nitrogen data layer, with insets by key coral reef regions. Values are distributed from 0 (lowest value in data layer) to 1 (highest value in data layer).

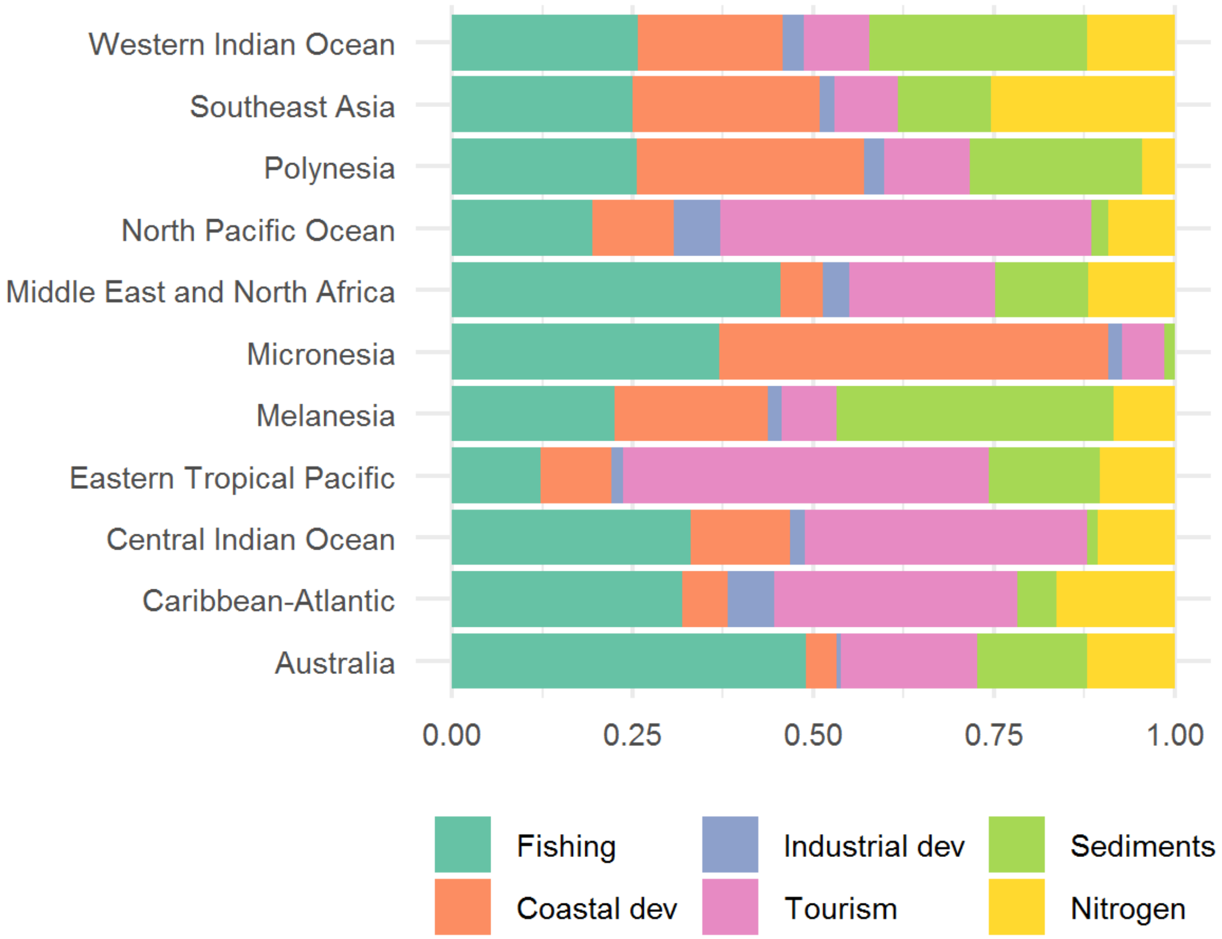


**Figure S7.** Regions vary in the relative proportion of top-ranked local pressures. Colours indicate pressure (see legend), and the length of each bar indicates the relative number of reef pixels within each region classified to each top pressure.

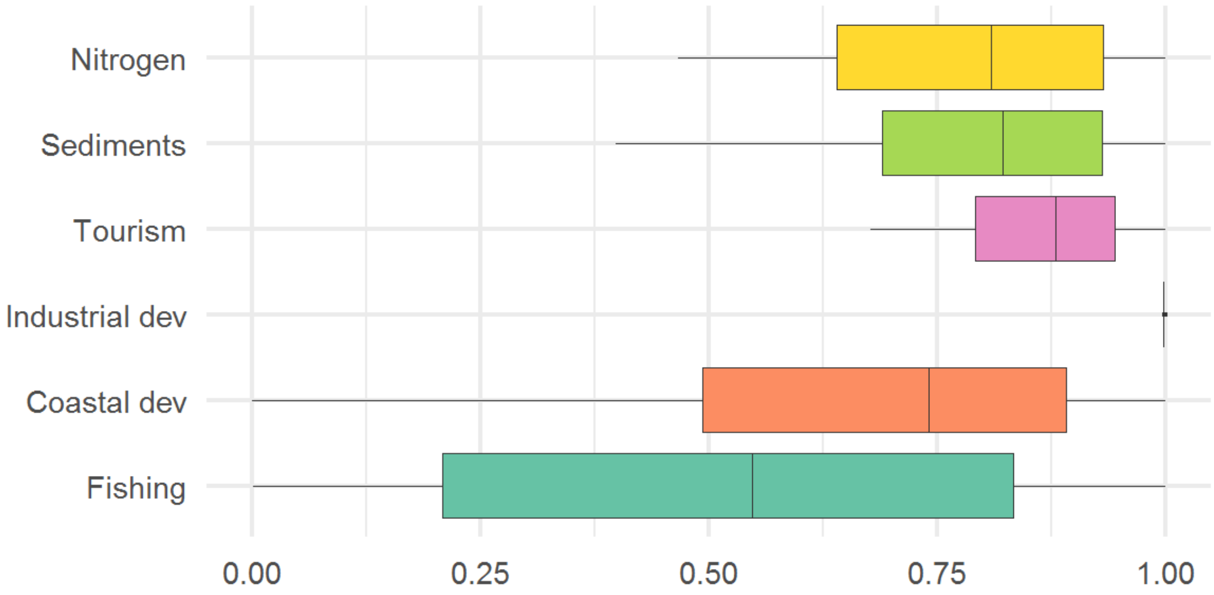


**Figure S8.** Pressure intensity of the top ranked pressures. Boxplots show the pressure percentiles when each pressure was identified as the top pressure in a reef pixel. When ranked as top pressures, nitrogen, sediments, tourism and industrial development have a higher pressure intensity, on average, than coastal development or fishing. Boxplots show the 25th, 50th (median) and 75th quantile, and whiskers are 1.5 * interquartile range (the distance between the first and third quartiles).

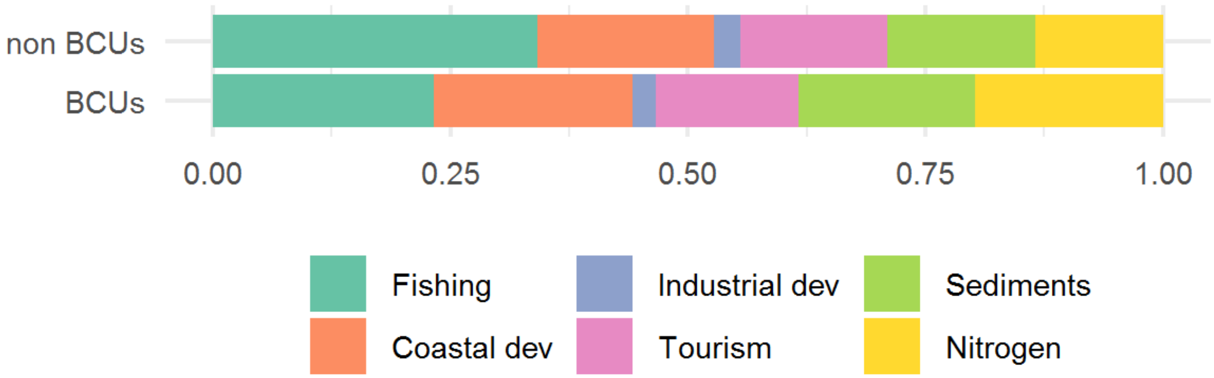


**Figure S9.** Comparison of top pressures between the 50 Reefs Bioclimatic Units (BCUs) and reefs outside potential climate refugia (non BCUs). Non BCUs typically have more fishing top pressures and fewer nitrogen pollution than BCU locations, but broadly the top-ranked pressures are very similar among BCU and non-BCU reef pixels.

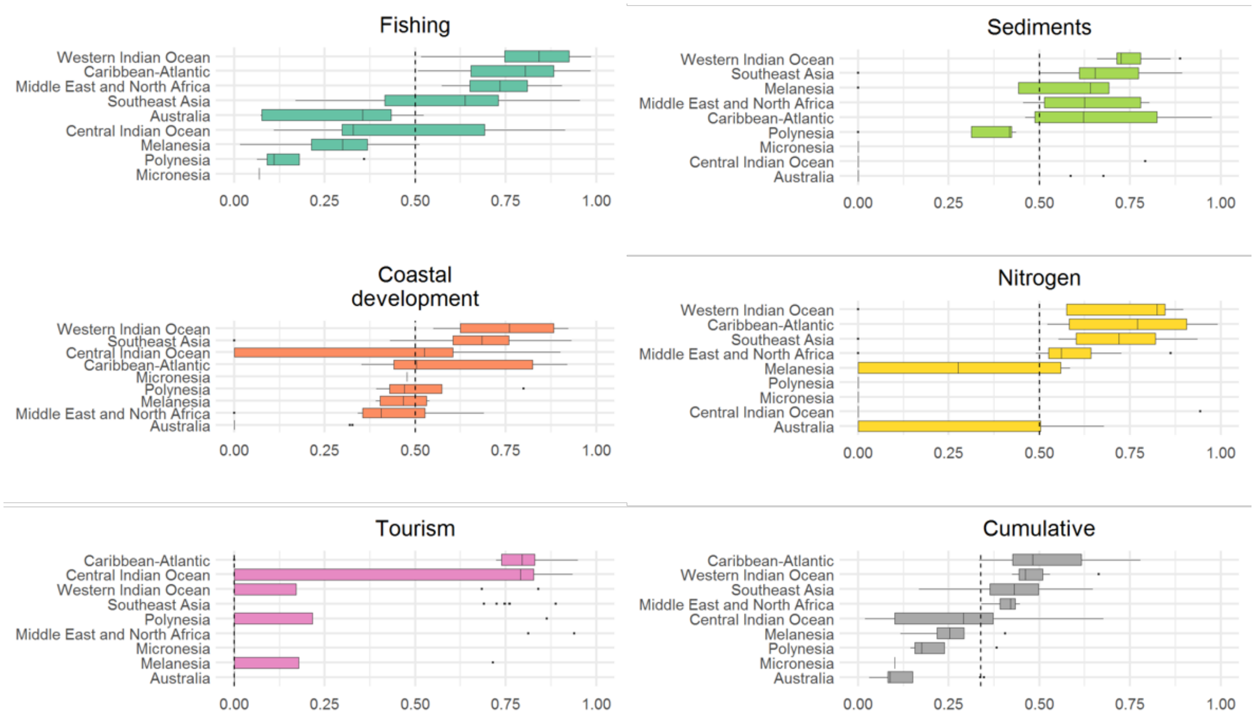


**Figure S10.** Distribution of the median impact scores of BCUs ($n$ = 83) by region. The vertical dashed line is the global median on all coral reef cells, regardless of whether they are included or not in a BCU.

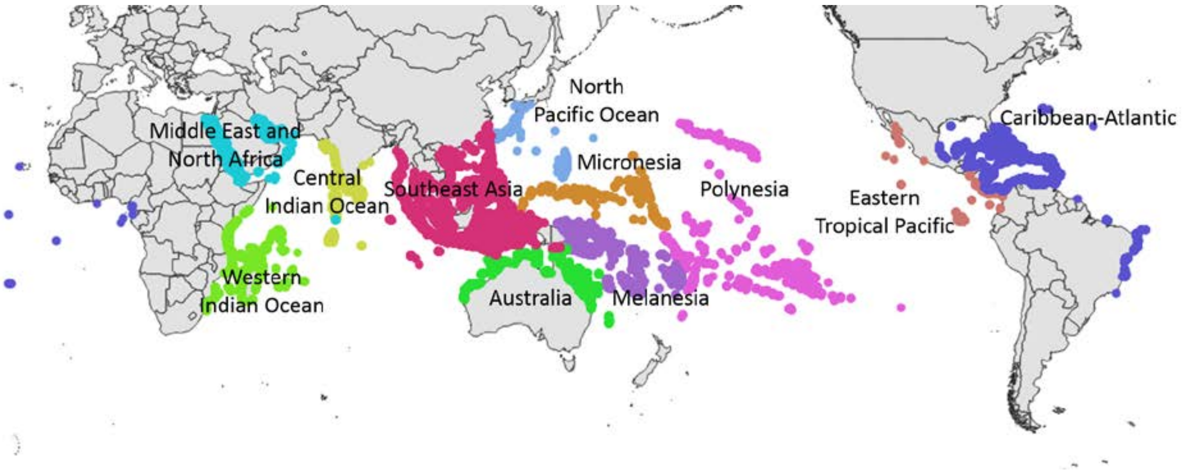


**Figure S11.** Coral reef regions used in this analysis. Different colours indicate different regions, and are broadly consistent with regions used by coral reef scientists and practitioners.

| | | | | | | |
|---|---|---|---|---|---|---|
| 1 km | 0.85 | 0.78 | 0.73 | 0.69 | 0.57 | 0.50 |
| | 5 km | 0.94 | 0.89 | 0.85 | 0.72 | 0.63 |
| | | 10 km | 0.97 | 0.94 | 0.81 | 0.70 |
| | | | 15 km | 0.98 | 0.86 | 0.75 |
| | | | | 20 km | 0.90 | 0.79 |
| | | | | | 50 km | 0.92 |
| | | | | | | 100 km |

**Figure S12.** Sensitivity of coastal development pressure to the distance threshold used to compute population counts. Values are Spearman's correlation coefficients among the percentiles of the coastal development pressure calculated using different radii. The main analysis uses a 5 km radius.
